# Towards rainy high Arctic winters: how ice-encasement impacts tundra plant phenology, productivity and reproduction

**DOI:** 10.1101/2021.06.10.447955

**Authors:** Mathilde Le Moullec, Anna-Lena Hendel, Matteo Petit Bon, Ingibjörg Svala Jónsdóttir, Øystein Varpe, René van der Wal, Larissa Teresa Beumer, Kate Layton-Matthews, Ketil Isaksen, Brage Bremset Hansen

## Abstract

The Arctic is warming rapidly, with winters warming up to seven times as fast as summers in some regions. Warm spells in winter lead to more frequent extreme rain-on-snow events that alter snowpack conditions and can encapsulate tundra vegetation in ‘basal ice’ (‘icing’) for several months. However, tundra climate change studies have mainly focused on summer warming. Here, we investigate icing effects on vascular plant phenology, productivity, and reproduction in a pioneer field experiment in high Arctic Svalbard, simulating rain-on-snow and resultant icing in five consecutive winters, assessing vascular plant responses throughout each subsequent growing season. We also tested whether icing responses were modified by experimentally increased summer temperatures. Icing alone delayed early phenology of the dominant shrub, *Salix polaris*, but with evidence for a ‘catch-up’ (through shortened developmental phases and increased community-level primary production) later in the growing season. This compensatory response occurred at the expense of delayed seed maturation and reduced community-level inflorescence production. Both the phenological delay and allocation trade-offs were associated with icing-induced lags in spring thawing and warming of the soil, crucial to regulating plant nutrient availability and acquisition. Experimental summer warming modified icing effects by advancing and accelerating plant phenology (leaf and seed development), thus increasing primary productivity already early in the growing season, and partially offsetting negative icing effects on reproduction. Thus, winter and summer warming must be considered simultaneously to predict tundra plant climate change responses. Our findings demonstrate that winter warm spells can shape high Arctic plant communities to a similar level as summer warming. However, the absence of accumulated effects over the years reveals an overall resistant community which contrasts with earlier studies documenting major die-off. As rain-on-snow events will be rule rather than exception in most Arctic regions, we call for similar experiments in coordinated circumpolar monitoring programmes across tundra plant communities.

## 1. Introduction

Global warming comes with stronger and more frequent extreme climate events, prompting research activity aimed at determining their ecological and evolutionary consequences for terrestrial biota (Grant et al., 2017; Harris et al., 2018; IPCC, 2021; Robinson 2022; Trisos et al., 2020). In the Arctic, temperatures are rising three times as fast as the global average and the particularly pronounced warming trends during winter have resulted in increasingly frequent and widespread winter warm spells (AMAP, 2021; Bintanja & Andry, 2017; Graham et al., 2017; Vikhamar-Schuler et al., 2016; You et al., 2021). Despite evidence for possible severe consequences of these warm spells on vegetation and carbon balance (Bjerke et al., 2017; Bokhorst et al. 2008, 2009, 2011, 2012, 2022; Phoenix & Bjerke, 2016; Treharne et al., 2019), studies have almost exclusively focused on the effects of rising summer temperatures and changes in snow depth (Bjorkman et al., 2020; Collins et al., 2021; Elmendorf et al., 2012; Prevéy et al., 2019; Rixen et al., 2022; Wipf and Rixen, 2010). Recently, studies comparing winter to summer warming effects on plant traits stress the neglected importance of winter temperatures for fine-scale regional patterns and traits sensitive to cold involved in resource acquisition and conservation (Bjorkman et al., 2018; Niittynen et al., 2020). Given the key role that tundra ecosystems play for global carbon cycling and climate feedbacks (Chapin et al., 2005; Zhang et al., 2020), it is now urgent to better understand how the addition of extreme warm spells outside the growing season shape Arctic plant communities.

Warm spells in winter often come with “rain-on-snow” events, i.e., when precipitation falls as rain instead of snow, often transforming the entire snowpack (AMAP, 2021; Bokhorst et al., 2016; Pan et al., 2018; Rasmus et al., 2018). Under such mild conditions, the snowpack can melt completely, exposing plants to thaw-freeze cycles that may directly cause vegetation damage (Bokhorst et al., 2009, 2011). Alternatively, and especially common in the high Arctic, rain and resulting meltwater can collect and freeze on the ground, releasing latent heath transferred to the soil and forms a layer of basal ice (hereafter referred to as ‘icing’) up to several decimetres thick (Peeters et al., 2019; Putkonen & Roe, 2003). Low-growing Arctic vegetation can remain entirely encapsulated in this ice layer for several months, until the onset of spring. Icing has the potential for ecosystem-wide consequences by directly or indirectly affecting several trophic levels, e.g. by reducing soil arthropod abundances (Coulson et al., 2000), disrupting snow conditions for small mammals’ survival and reproduction (Kausrud et al., 2008), and causing starvation-induced die-offs in large herbivores, which in turn can influence predators and scavengers (Hansen et al., 2013; Sokolov et al., 2016). However, the immediate and lasting effects of icing on high Arctic plants’ productivity, reproduction, and phenology – fuelling the tundra food web – remain largely unknown.

The increased occurrence of regional Arctic vegetation ‘browning’ (i.e., decrease in expected ‘greening’ under warmer climate) has been associated to several factors, including winter warm spells associated to thaw-freeze cycles (Bokhorst et al., 2008, 2009; Phoenix & Bjerke, 2016), but also to icing (Bjerke et al., 2017, Milner et al., 2016, Vickers et al., 2016). This contrasts with the large-scale ‘Arctic greening’ pattern associated to shrub expansion and improved plant growing conditions (Epstein et al., 2012; Frost et al., 2019; Myers-Smith et al., 2011; Walker et al., 2006). However, changes in winter conditions can also exacerbate the effect of summer warming alone (Frei and Henry, 2021), thus adding complexity and spatial variability for vegetation productivity trends (Berner et al., 2020; Frost et al., 2019; Kelsey et al., 2021; Myers-Smith et al., 2020), pressing needs for teasing appart effects of icing.

Physiologically, icing can lead to plant cell death, either through frost damage or stress-induced anoxia metabolite accumulation from anaerobic respiration (Bokhorst et al., 2010; Crawford et al., 1994; Preece & Phoenix, 2014). Studies investigating the effects of icing described that evergreen shrubs and lichens were negatively affected, either through altered leaf physiology and/or shoot and individual mortality (Bjerke, 2011, 2017, 2018a, 2018b; Milner et al., 2016; Preece et al., 2012). In comparison, deciduous shrubs and graminoids showed higher tolerance, yet, there was still evidence of damage and biomass reduction (Bjerke et al., 2018b; Preece & Phoenix, 2014). In addition, icing can indirectly affect the vegetation by altering soil thermal properties (Putkonen & Roe, 2003), delaying plant phenology as seen under late snow-melt (Assmann et al., 2019; Frei and Henry, 2021; Semenchuk et al., 2016), or alternatively, icing may protect plants against winter herbivory.

Plants facing sub-optimal conditions may channel more energy into either growth or reproduction, modifying existing energy allocation trade-offs (Bazzaz et al., 1987; Jónsdóttir, 2011). Induced stress can enhance vegetative growth, thereby constraining or delaying other reproductive responses, i.e., flower phenology and/or number (Chapin, 1991; Orcutt & Nilsen, 2000). Such responses are found in Arctic plants coping with shortened growing seasons (e.g. late snowmelt, Cooper et al., 2011; Wipf & Rixen, 2010), damages from herbivory (Jónsdóttir, 1991; Skarpe & Van der Wal, 2002), thaw-freeze events (Bokhorst et al., 2011), as well as icing (Milner et al., 2016; Preece et al., 2012). In these latter studies simulating icing, Milner et al. (2016) showed high mortality of apical meristems and entire shoots in the evergreen shrub *Cassiope tetragona* in the high Arctic, but increased growth of surviving shoots, at the cost of reduced flower production (Milner et al., 2016). Along these lines, Preece et al. (2012) expected delayed leaf phenology following icing in the low Arctic, but instead found accelerated leaf emergence in spring in *Vaccinium myrtillus* and *V. vitis-idaea*, suggesting a reallocation of resources to the remaining living shoots, yet, the cost of flowering was unclear. Species like *Empetrum nigrum* were resistante to icing (Preece et al., 2012).

Warmer summers may counteract expected negative effects of icing through greater activation of growth hormones fluxes and enhanced nutrient cycling and availability (Chapin, 1991; Frei and Henry, 2021; Gornall et al., 2011; Sundberg et al., 2000). Again, responses may vary across species and tissue types, i.e., if the tissues are directly or indirectly exposed to icing in addition to summer warming. For instance, high Arctic perennial species commonly produce flower buds the previous summer (Arft et al., 1999, Barrett and Hollister 2016), thus directly experiencing icing conditions, while vegetative tissues grow generally in direct response to June-July temperatures (Van der Wal & Stien, 2014). Possible delay in phenology under icing could be altered by summer warming, advancing more reproductive than vegetative, as well as early than late phenophases (Collins et al., 2021), with greater effects on early than late flowering species (Prevéy et al. 2019). However, regardless of summer warming, the duration of phenophases tend to be fix (Barrett and Hollister 2016; Collins et al., 2021; Semenchuk et al., 2016).

Plot-level field experiments are powerful tools to investigate fine-scale plant response to simulated climate change scenarios (Bjorkman et al., 2020; Bjorkman et al., 2018; Elmendorf et al., 2015). Here, we assessed how experimental winter rain-on-snow events and subsequent icing affect productivity, reproduction, and phenology of a high Arctic plant community. To achieve this, we simulated plot-level rain-on-snow and ice formation in winter over five consecutive years. Using a full factorial design, we also increased summer temperatures via open top chambers, following the protocol of the Arctic and Alpine International Tundra Experiment (ITEX, Henry & Molau, 1997). During each subsequent growing season, we determined plant species and community responses to treatments. We expected icing to cause overall negative effects, i.e., to 1) delay the plants’ phenology due to a delaying effect of icing on thawing processes; 2) reduce the overall plant productivity (‘browning’); and 3) reduce the flower production due to re-allocation of resources to growth. We anticipated the addition of summer warming to overall reduce negative effects of icing, while the net effects are difficult to predict *a priori*, given the scarcity of research in this field, the many mechanisms involved, and the likely variation across species and tissue types.

## 2. Methods

### Study site

The high Arctic Svalbard (78°17’N, 16°02’E) is a region warming twice faster than the Arctic average and five-seven times faster than the global average (Isaksen et al. 2022b). The experiment was located in the valley of Adventdalen and lasted for five consecutive years (January 2016–August 2020). Near the study site, the annual mean air temperature recorded at Svalbard Airport (15 km away) was −5.9 °C for the reference period 1971-2000 (Hanssen-Bauer et al., 2019), and −2.2 °C for the study period. Local winter (December–February) temperature rise five to seven times faster than in summer (June–August; Isaksen et al., 2022), thus at a greater rate than at the circumarctic scale. The interior fjord and valley areas around Adventdalen are some of the driest in Svalbard, with an average annual precipitation of 196 mm for the reference period (Hanssen-Bauer et al., 2019) and 223 mm for the study period. In comparison to the reference period, median RCP8.5 projections for Svalbard anticipate an increase in annual temperature and precipitation of about 7-9 °C and 20-40%, respectively, towards 2100 (Hanssen-Bauer et al., 2019). The frequency of rain-on-snow events has increased dramatically since a regime shift around the year 2000 (Hansen et al 2014, Peters et al. 2019), and Svalbard currently experience conditions soon expected across large parts of the Arctic (AMAP, 2021; Bintanja & Andry, 2017).

Soil thermal conditions on Svalbard are linked to the underlying permafrost which warms at a rate of 1.4 °C per decade at the permafrost table (2 m depth), observed 6 km away from our study site between 2002–2018 (Etzelmüller et al., 2020). At the study site, the sub-surface (5 cm depth) soil temperature was on average 4.7 °C in June and 7.5 °C in July (see methodology below, Table S1a), the two most important months for plant growth (Van der Wal & Stien, 2014). The volumetric water content (i.e., soil moisture) is relatively low for mesic communities, drying throughout the summer season from about 45% in early June to 25% mid-August at 10 cm depth (Table S1a). Soil moisture and sub-surface temperature are strongly negatively related (Table S1b).

The tundra plant growing season starts immediately after the soil-thaw onset (Descals et al., 2020). Adventdalen, which is in the bioclimatic subzone C ‘middle Arctic tundra’ (CAVM Team, 2003), has a growing season of approximately three months, starting in mid-late May (Table 1). The plant community studied here was a mesic grass and moss-rich dwarf shrub heath, dominated by the prostrate deciduous dwarf shrub *Salix polaris* and the grass *Alopecurus borealis*. Other abundant vascular plants were the horsetail *Equisetum arvense*, the rush *Luzula confusa*, the forb *Bistorta vivipara* and the grass *Poa arctica* (Fig. S1). The most abundant bryophytes were *Sanionia uncinata, Tomentypnum nitens* and *Polytrichastrum* spp. Nomenclature follows http://panarcticflora.org/ for vascular plants and Frisvoll and Elvebakk (1996) for bryophytes.

**Table 1.**
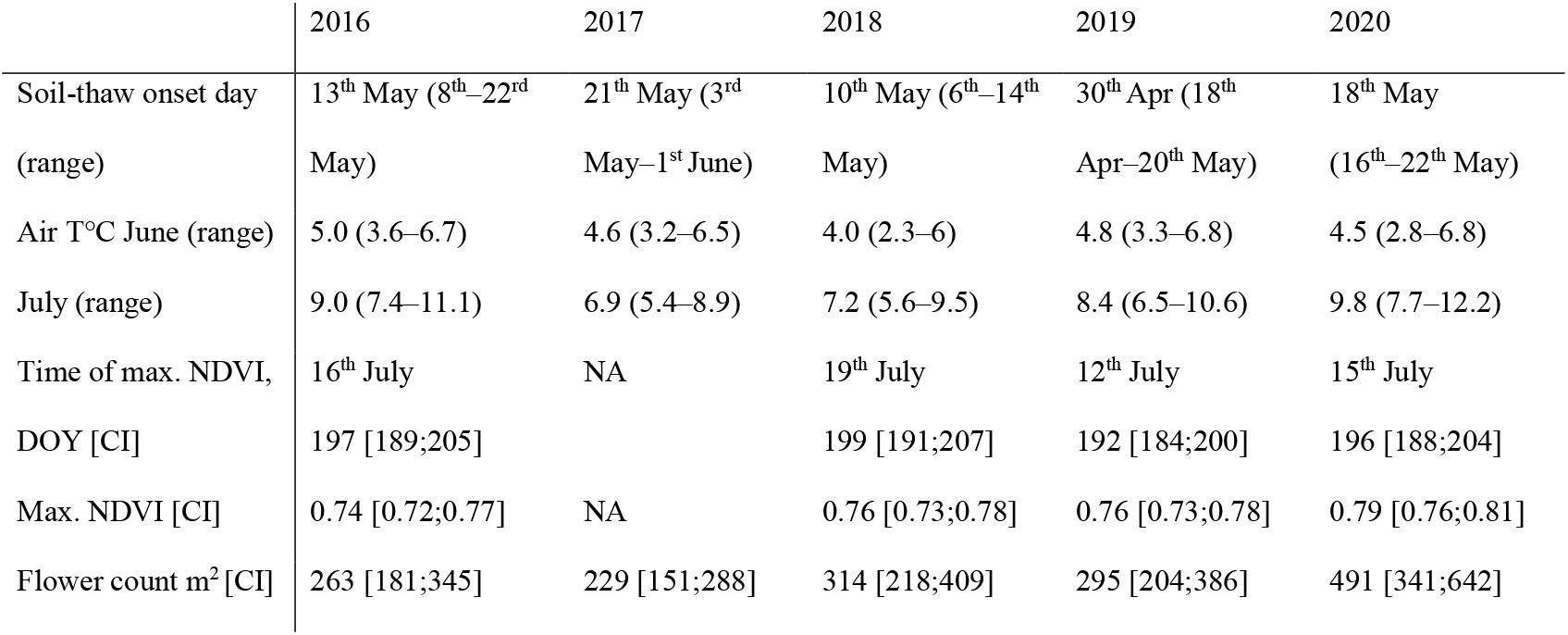
Overview of annual mean biotic and abiotic measurements. The daily average air temperatures were recorded at the Svalbard Airport (available on https://seklima.met.no/), while the soil-thaw onset day, maximum Normalized Difference Vegetation Index (max. NDVI) metrics, and the community inflorescence counts (Flower count) were recorded in control plots. Predicted means are presented with their 95% confidence intervals (CI), or range of values. The ‘time of max. NDVI’ is interpreted as the peak growing season. DOY = Day-of-year (i.e. Julian day, where 1 = 1 January).

### Experimental design

The study design followed a full-factorial randomized block design with four treatment combinations (2 levels of icing × 2 levels of warming), replicated three times within each of the three blocks (Fig. 1). In summer 2015, the three blocks (150–780m apart) were selected in homogenous mesic tundra. Within each block, we chose 12 plots of 60 × 60 cm with similar communities (n = 36 plots, Fig. 1). The four treatments, icing (*I*), summer warming (*W*), a combined winter icing and summer warming treatment (*IW*), and control (*C*), were randomly assigned to plots. Plots received the same treatment each year, starting in winter 2015–2016 (Table S2). This allowed us to investigate both short-term (one year) as well as longer-term (up to five years) plant responses. However, not all traits and parameters were measured in all years, hence some results are presented based only on a subset of years.

**Fig. 1.**
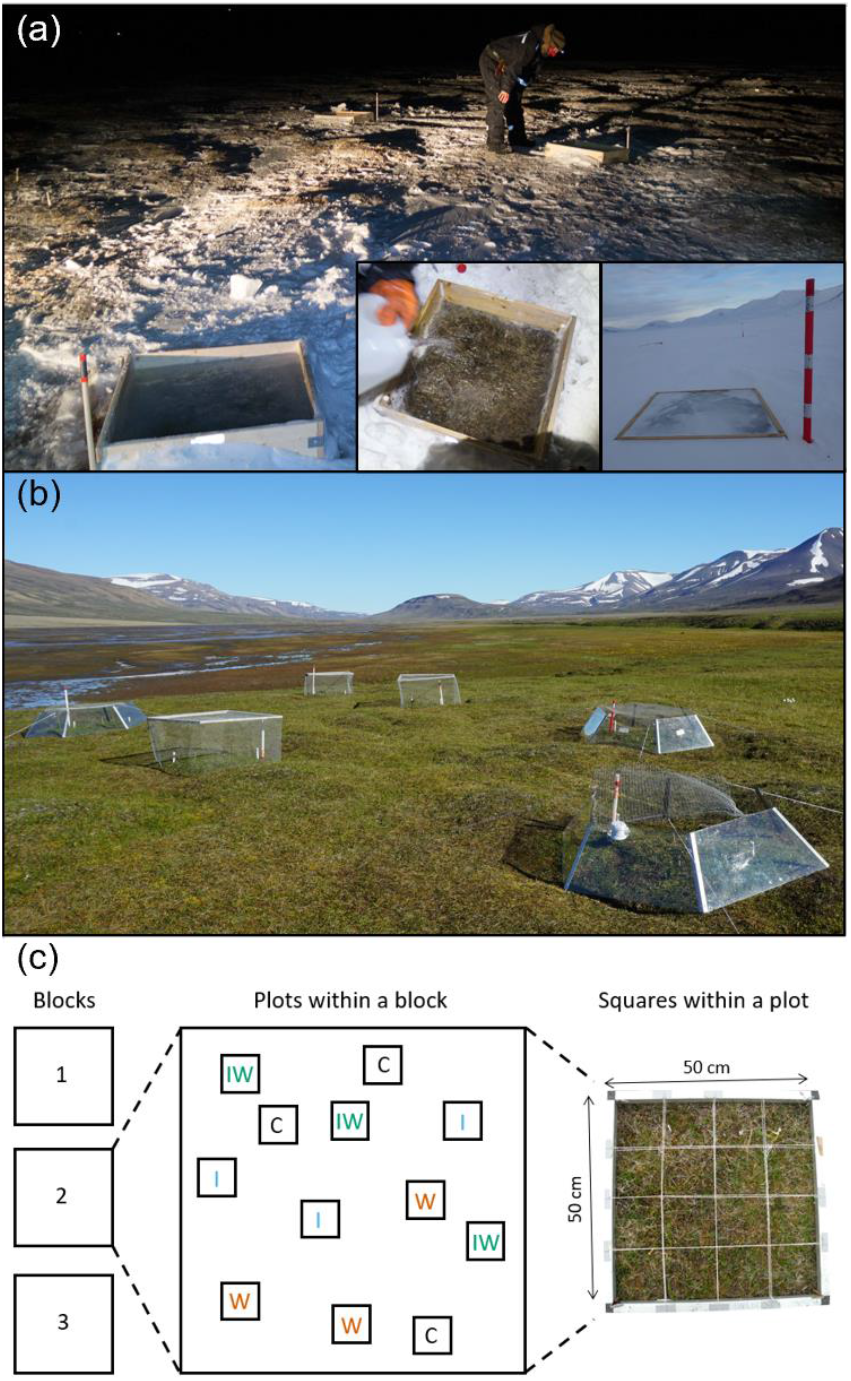
The experimental set up in the Adventdalen valley, Svalbard. Overview of the field site (a) in the polar night, when applying the icing treatment by filling a wooden frame with water, and (b) in summer with the open top chambers (plexiglass hexagons) simulating warming and net cages to protect against herbivory. (c) Experimental full-factorial randomized block design with the vegetation frame fitting the plot size. C = control, I = icing, IW = icing × warming, W = warming. Picture credits: Ø. Varpe and M. Le Moullec.

#### Winter icing

The experimentally formed ice was applied to *I* and *IW* plots within a period of 2–3 days in January–February (Fig. 1a). Snow was carefully removed from all plots to apply the same level of disturbance, and any occurrence of natural ice was recorded. Natural ice was particularly common in 2017, with partial ice coverage occurring in all plots (3 cm average thickness). Snow was immediately placed back on non-icing (*C* and *W*) plots, while for *I* and *IW*, a 13 cm high wooden frame (60 × 60 cm) was placed around the plot and gradually filled up with cold water (for 1-3 days) until it was full of solid ice (Fig. 1a), following Milner et al. (2016). Frames had no influence on the snow accumulation pattern later in winter and they were removed at the beginning of snowmelt. Snow- and ice-melt occurred almost simultaneously during spring meltwater floods.

#### Summer warming

Right after snowmelt (Table S2), hexagonal open top chambers (1.4 × 1.4 m basal diagonal) were deployed over the *W* and *IW* plots following the specifications given by the ITEX protocol (Henry & Molau, 1997). To avoid confounding factors, all plots were excluded from grazing herbivores during the snow-free season using metal net cages (*I* and *C* plots) or nets on top of the open top chambers (*W* and *IW* plots; mesh-size 1.9 cm × 1.9 cm, Fig. 1b). Open top chambers and nets were all deployed and removed on the same day in the start and end of the growing season.

### Field measurements

Soil temperatures were recorded with iButton loggers (type DS1921G-F5, ± 1.0 °C accuracy, 0.5 °C resolution) every 120 minutes in summer and 240 minutes in winter. Loggers were placed in all plots at depths of both 2 cm and 5 cm. In 2018, we also recorded temperatures at 10 cm and 20 cm depth (not in *IW*). We estimated the onset of soil-thaw in spring for each plot based on logger data at 5 cm depth; if soil sub-surface temperatures were ≥ 0 °C for a minimum of 10 consecutive days, the first day of this period defined the soil-thaw onset. We also monitored surface air temperature (5 cm high) with HOBO loggers (type U23-003/UA-001; ± 0.2°C accuracy) in a limited number of plots in *C* and *W* from 15^th^ of June to 1^st^ of September 2016-2018, every 30 min (Table S1a).

To assess treatment effects on plant production, we monitored the vegetation weekly during the growing season by measuring Normalized Difference Vegetation Index (NDVI, a proxy for plant productivity with a value from −1 to 1; Pettorelli et al., 2005). This was done at the central part of each plot (30 cm diameter) with a Skye SpectroSense2+ handheld device. NDVI typically increases in the first part of the growing season until reaching a maximum, to then drop progressively in the second part of the growing season. We considered the peak growing season to be the average day *C* plots reached the maximum NDVI (Table 1). As an additional measure of plant productivity, we measured community-level vascular plant species abundance (hereafter referred to as ‘relative abundance’) in the period after the peak growing season (end of July, Table 1), in 2016–2019 (Table S2). To estimate relative abundance, we used the point intercept methodology (Bråthen & Hagberg, 2004), with a 50 × 50 cm frame elevated above the canopy (~20 cm high) and with 25 evenly distributed points. The points were marked by crossings of double strings with the frame to give a 90° projection. At each point, a wooden pin of 3 mm diameter was lowered down onto the moss layer and all ‘hits’ of vascular plants were recorded.

To further study vegetative responses of the dominant vascular species, we collected *S. polaris* leaves (before leaf senescence, Table S2) to measure leaf area, leaf dry weight and specific leaf area (SLA, the ratio of leaf area to dry mass). The protocol and sample size of leaf collection varied between years (Table S2), preventing direct comparison of leaf-size traits between years, but still enabling within year comparisons of *I, IW* and *W* effect sizes relative to *C*. Leaves were kept moist after collection until scanned. Leaf area was measured in imageJ 1.48 (Schindelin et al., 2015). Subsequently, leaves were oven-dried at 60 °C for four days and weighed on an XS204 METTLER TOLEDO scale to the nearest 0.01 mg.

To assess treatment effects on flowering frequency, the total number of inflorescences were counted for the species *S. polaris* (males and females [catkins]), *B. vivipara, A. borealis, L. confusa and P. arctica* in mid-July each year (Table S2). We counted inforescences in a 50 × 50 cm frame (Fig. 1c), subdivided into 16 sub-squares.

To assess treatment effects on the phenology of the dominant vascular plant, we measured the most advanced vegetative and reproductive phenological stages (‘phenophases’) of *S. polaris* in each of 16 sub-squares per plot of the 50 × 50 cm frame. We used the following vegetative phenophases: 1) leaves starting to unfurl, 2) leaves fully expanded, 3) start of senescence, and 4) leaves fully senesced. The reproductive phenophases were 1) distinct inflorescence buds visible, 2) buds recognisable as female or male, 3) receptive stigmas (female) and open anther releasing pollen (male), 4) stigma and anthers withered, and 5) seed dispersal. The highest number of phenological monitoring rounds took place in 2018, whereas no phenological monitoring was done in 2020 (Table S2).

### Statistical analysis

We derived different metrics from the NDVI data by fitting a generalized additive model (GAM) to each plot for each year, using the repeated measurements over a growing season. We employed the function ‘gam’ from the mgvc package (Wood et al., 2015) in software R-3.6.3 (R Core team, 2020), fitted with restricted maximum likelihood. Model fit was good (visual interpretation and k-indices > 1), except for summer 2017 when measurements stopped before the peak growing season, i.e., they were still in the increasing phase (Table S2). Thus, we removed 2017 from all NDVI analyses. For all other years, we predicted daily NDVI values for each plot and derived 1) the maximum NDVI value, and 2) the day-of-year when the maximum NDVI was reached. We also computed 3) the cumulative NDVI in the first part of the growing season (hereafter ‘cumulative start’) by summing daily values from the first day of measurements to the day when *C* plot, on average, reached the maximum NDVI, 4) the cumulative NDVI in the second part of the growing season (hereafter ‘cumulative end’) by summing daily values from the day when *C* plot, on average, reached the maximum NDVI to the last measurement day of the season, and 5) the cumulative NDVI across the growing season (hereafter ‘cumulative total’) by summing daily values from the first to the last day of measurements. These cumulative NDVI metrics were then standardized by the number of days over which the integral was computed. This was necessary because the field season length differed between years, but not between treatments within year.

We used (generalized) linear mixed-effect models as the main tool to analyse our data and account for the hierarchical spatial and temporal structures of our study design, using the functions ‘lmer’ and ‘glmer’ from the lme4 R package (Bates et al., 2015). We built a separate model for each response variable, i.e., soil temperature, onset of soil-thaw, NDVI (different metrics), plant relative abundance, leaf size traits, inflorescence number, and phenophases. We conducted four different models for each response variable. 1) We computed the treatment effect sizes or predicted mean over years (treatment [category] was set as fixed effect) and the temporal replication was accounted for by including year as a random intercept, or day-of-year nested within year when measurements were also replicated within a given year (e.g., NDVI). 2) We calculated the annual effect sizes or predicted means (‘treatment [category] × year [factor]’ interaction as fixed effects). 3) We tested for an overall icing-warming interaction effect over years, (icing [0,1] × warming [0,1] interaction as fixed effects) with ‘year’ or ‘day-of-year’ nested within ‘year’ as random intercepts. 4) We tested for an annual icing-warming interaction effect, (icing [0,1] × warming [0,1] × ‘year [factor] interaction as fixed effects. For all models, the random intercept structure (i.e., variation in means between replicated units) always included the nested plot (n = 36) structure within blocks (n = 3). It also included sub-squares nested within plots (16 sub-squares per plot, n = 576) for measurements of leaf traits (in 2019 and 2020, Table S2), inflorescence count, and phenology. In 2018, leaf traits included information on leaves nested within shoots (Table S2). Statistical significance can be interpreted as when the 95% confidence interval (CI) of estimates did not overlap the predicted mean of *C*.

Data from air and soil temperature and NDVI metrics were best summarized by a normal distribution and count data from plant relative abundance and inflorescence counts by a Poisson distribution. Leaf size traits were log-transformed to follow a normal distribution. Each phenophase was converted to a binary data format to fit a logistic regression from a binomial distribution. Percentage change reported were calculated for back-transformed predicted means, in comparison to controls.

## 3. Results

### Soil temperature and soil-thaw onset

The liquid water added during the simulated rain-on-snow on both icing (*I*) and icing × warming (*IW*) plots released latent heat, increasing sub-surface soil temperatures for three days, by on average 2.9 °C [2.3;3.5] (results are reported as mean effect sizes [95% CI] in comparison to controls [*C*] over 2016–2020, unless specified differently) (Fig. S2).

During spring snowmelt, sub-surface soil-thaw onset was delayed by 6.0 [2.2;9.9] days in *I* and 4.4 [0.4;8.5] days in IW, while it remained unaffected in warming (*W*) (−0.9 [−5.0;3.1] days, Fig. 2a). However, the sub-surface soil-thaw onset, and the period over which it occurred, varied considerably between years (Table 1). For instance, the thawing onset of *C* occurred on average on 30^th^ of April in 2019, ranging over a period of 32 days, while it occurred on the 21^st^ of Mai over a period of 6 days in 2020 (Table 1). At 20 cm depth (data only from 2018), soil-thaw onset was delayed by 9.0 [1.3;16.7] days in *I*, after staying at a constant temperature of −0.5 °C for about two weeks (i.e., ‘the zero curtain period’, Outcalt et al., 1990; Fig. 2b), resulting in lagged soil temperature increases in *I* (i.e., significantly lower during entire June, Fig. 2b).

**Fig. 2.**
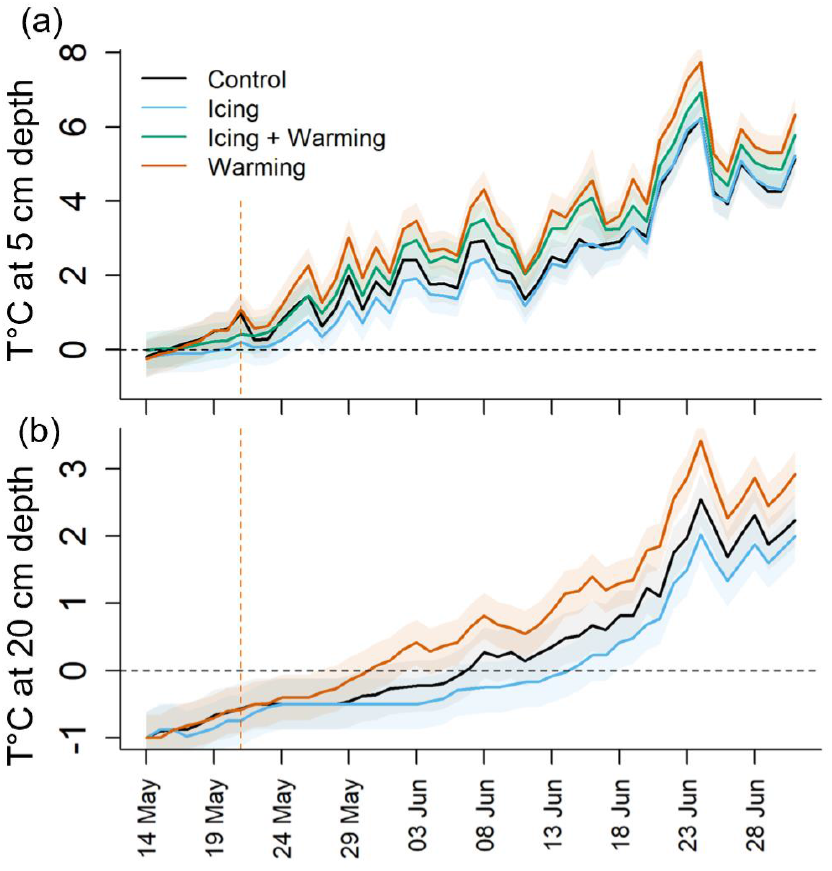
Daily estimates of soil temperatures during the soil-thawing period from mid-May to end of June 2018, at (a) 5 cm depth (i.e., sub-surface), and (b) 20 cm depth. Shaded areas represent 95% CIs and the orange vertical dashed lines shows the day the warming treatments started.

The presence of open top chambers increased soil sub-surface temperatures by 0.9 [0.7;1.2] °C in *W* and 0.8 [0.6;1.1] in *IW* (at 2 cm depth in June-August, over 2016, 2018 and 2019) and surface air temperature (5 cm high, 2016-2018) by 0.8 [0.7;0.9] °C in *W* (Table S1a). The soil volumetric water content did not vary between treatments but varied between years (Table S1a). The first and last year of the experiment were the warmest summers (2016 and 2020), while 2017 was the coldest (Table 1). Soil was the wettest in 2020 and the driest in 2019 (Table S1).

### Vegetative responses: NDVI metrics, relative abundance, leaf size traits and phenology

Right after soil-thaw onset, the NDVI curves increased rapidly until reaching a maximum in mid-July in *C* (15^th^ of July on average, Table 1 and Fig. 3a) and decreased slowly thereafter (Fig. S3). The maximum NDVI timing was delayed by on average 4 [2;7] days in *I*, and consistently so across years (Fig. 3i). This delay was reduced in *IW* in most years (2 [−1;4]) while it was −3 [−5;0] days earlier in *W* (Fig. 3i and Table S4). Maximum NDVI values themselves were higher in *I* and *IW* in the first year of treatment (2016), while this effect vanish in later years (Fig. 3j and Table S4). The last year of treatment (2020) had a high maximum NDVI compared to previous years (Fig. 3b and Table 1), where even *C* reached values similar to those in *I* and *IW* in other years (Fig. S3, Table S4). Maximum NDVI values were unaffected by *W* (Fig. 3j and Table S4).

**Fig. 3.**
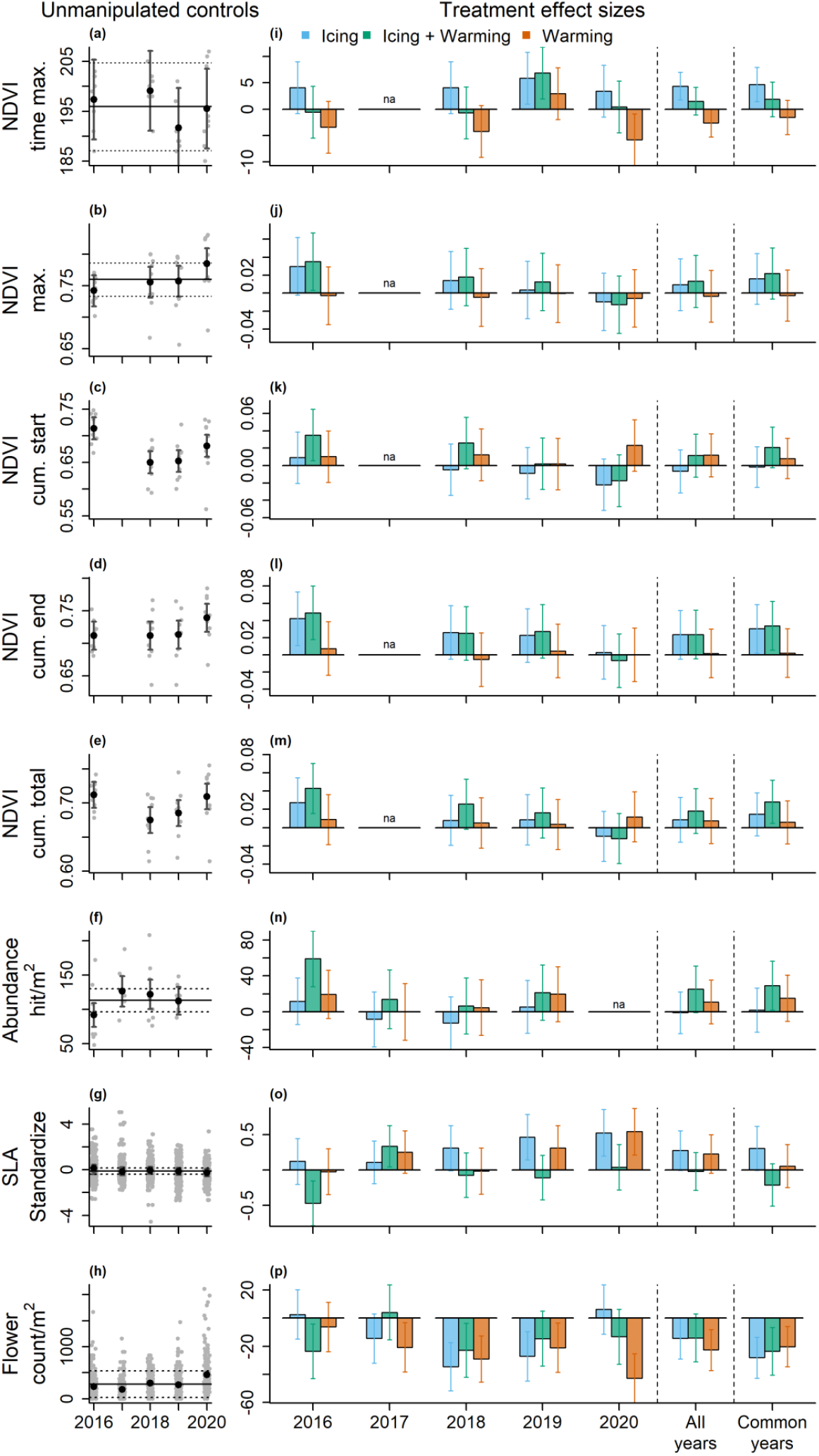
Effects of experimental treatments on several Normalized Difference Vegetation Index (NDVI) metrics, relative abundance (abundance) of vascular plants, specific leaf area (SLA) of *Salix polaris* and the community inflorescence count (flower count). (a-h) Model predictions (black dots) and their 95% confidence intervals (CIs) for unmanipulated control plots, separately for each year. The horizontal full and dashed lines represent the among-year model predictions and their 95% CIs, which are only displayed for variables comparable between years (i.e., with comparable sampling periods and/or design). Dots in the background show the raw data at the plot (n = 9) or sample replication within plot (n range [109-214]) level, which were jittered for display purposes. Model predictions and their CIs were back-transformed on the response scale prior presentation. (i-p) Effect sizes and their 95% CIs for the effect of treatments, displayed separately for each year (left panel - year is used as a fixed-effect in interaction with treatment in the model) and across all years (right panel - year is used as a random-effect in the model). In the left panel, the reference level at 0 effect size represents within-year unmanipulated controls (corresponding to the model predictions in a-h) and effect sizes refer to within-year treatment effects. In the right panel, the reference level at 0 effect size represents across-year unmanipulated controls (corresponding to the horizontal lines in a-h) and effect sizes refer to across-year treatment effects. Abbreviations as follows: NDVI Time max. = Time of maximum NDVI as day-of-year; NDVI max. = Maximum NDVI values from −1 to 1; NDVI cum. start = NDVI cumulative start. NDVI cum. end = NDVI cumulative end. NDVI cum. total = NDVI cumulative total (see Material and Methods for details).

The shape of the NDVI curves varied among treatments, particularly after the maximum NDVI. This effect was captured by the NDVI cumulative metrics. For the first part of the growing season, only *IW* increased the NDVI ‘cumulative start’, and remained high for the second part of the growing season (NDVI ‘cumulative end’), resulting in an overall increased NDVI (‘cumulative total’) across the growing season (Fig. 3k-m, Table S4). In the second part of the growing season, *I* caught-up with *IW*, and thus also exceeded the NDVI ‘cumulative end’. NDVI cumulative metrics of *W* did not differ (Fig. 3b). The treatment effect sizes of NDVI metrics decreased over the years (except for the timing of the maximum NDVI, Table S4), but still were of comparable magnitude as annual variation in *C* (Fig. 3).

Vascular plants’ relative abundance was unchanged under *I* (3% [−11;19] change, Fig. 3n and Table S5), with a slight tendency for higher abundance of *S. polaris* (21% [−17;75]). However, under *IW*, the relative abundance was on average 26% [9;46] higher, with large effects on the first treatment year (2016, 69% [43;99] change, Fig. 3n and Table S5). These patterns were largely consistent with the NDVI metric for the entire growing season (‘cumulative total’) and driven by the two most abundant vascular plants, the shrub *S. polaris* and the graminoid *A. borealis* (Fig. S4), which relative abundance had a large influence on the NDVI patterns (based on among-plot correlations, Table S3). *A. borealis* was even three times more abundant in *IW* on the first treatment year (Fig. S4 and Table S5). Under *W*, the relative abundance tended to increase by 13% [−2;31], a change not related to *S. polaris* which decreased by −32% [−54;-1] (Figs. S3 and Table S5). The vascular plant’ species community composition remained overall unchanged across the study period (Fig. S5).

The SLA of *S. polaris* was higher in *I*, and this difference significantly increased over the years (Fig. 3o and Table S8), being particularly pronounced from 2018 onwards. In those years (2018-2020) dry weight decreased more than the area, with a respective change of −19% [−28;-1] and −14% [−28;4] (Table S6). Although the leaf area and dry weight tended to be lower in all treatments and all years, their ratio (i.e., SLA) was not different in *IW* and *W*, with large inter-annual variation (Figs. 3o, S6 and Table S6).

No significant interaction effect was found between *I* and *W* in vegetative traits, except for SLA in 2020 and early leaf phenology (see below), i.e., there were no strong indications of an overall modifying effect of summer warming on the response to icing (Tables S4-S6).

### Reproductive response: inflorescence production

The total number of vascular plant inflorescences in this mesic community was significantly reduced across all treatments, and this particularly strongly over the years with all types of measurements (2016, 2018 and 2019), showing a reduction by one third (Fig. 3p and Table S7). Yet, *I* had no effect in the first and last treatment year (i.e. 2016 and 2020, Fig. 3p, Table S7). Although *S. polaris* produced the highest number of inflorescences (catkins and male flowers), and drove this overall pattern of annual variation, inflorescences of the forb *B. vivipara* and graminoids were severely impacted by *I* (and *IW*) in all years (respectively by −52% [−72;-27] and −67% [−95;-11] reduction, Fig. S7 and Table S7). Under *W*, *S. polaris* inflorescences were also strongly reduced (−70% [−81;-56]), while graminoid inflorescences were enhanced (146% [46;225]). Note that in 2020, there was a record number of inflorescences in the community, with more than twice the amount compared to any other study year (Fig. 3h, S7 and Table 1). For instance, even if the greatest reduction of inflorescences under *W*, occurred in 2020 (Fig. 3p), after a progressive decrease over time (Table S8), it was still the year with the greatest inflorescence production under this treatment (Fig. S7 and Table S7).

The interaction effect of *I* and *W* was positive, in other words, there were more inflorescences under *IW* than expected if the effects of *I* and *W* were additive (Table S7).

### Phenological responses

Early phenophases of *S. polaris* leaves developed later in *I*, while they developed earlier in *IW* and *W* (i.e., when 50% of the *C* reached the phenophase of unfurled leaf, 10% in *I* and 90% in *IW* and *W* did so, Fig. 5a). Hence, the unfurled stage was reached 4 [1;8] days later in *I* and −4 [−8;-1] days earlier in *IW* and *W* (Fig. S8). Despite this delay in *I*, the duration to unfold leaves (from unfurled to fully expanded) was shortened by one third compared to *C* and the other treatments (−3 [−4;-1] days shorter, Fig. 5a-b and Table S10). This resulted in leaves being fully expanded with no apparent delay under *I* (2 [−2;5] days), but −5 [−8;-2] days earlier in *IW* and *W* (i.e., when less than 10% of the *C* and *I* reached the fully expanded stage, all plots had already reached it *IW* and *W*, Figs. 5b, S8, S9, Table S9 and S10). However, the onset of leaf senescence happened almost simultaneously across treatments, and this pattern occurred consistently across years (Figs. 5c-d, S8 and S9). Yet, this timing of leave senescence varied among years with more than 10 days in *C* (Fig. S8a). The duration of senesced was unchanged in *I*, while in *IW* and *W* it was 30% faster (−2 [−3;-1] days shorter). Note that in contrast, *IW* and *W* leaves stayed 20% longer time fully expanded (6 [2;8] days; Table S10).

Several *S. polaris* reproductive phenophases were reached later in *I* but earlier in *IW* and *W* compared to *C* and these differences persisted throughout years, being particularly pronounced in 2017 (Figs. 5 and S8b-c). The difference in timing was largest during seed dispersal. In 2018, it occurred slightly later in *I* (5 [−2;12] days), but −11 [−17;-6] and −5 [−12;2] days earlier in I*W* and *W* respectively (i.e., when 50% on the *C* reach seed dispersal, 19% in *I*, 96% in *IW* and 86% in *W* had reached it; Figs. 5 and S9). The ealrier seed dispersal in *IW* coincided with a 30% shorter duration for seeds to mature (−8 [−14;-1] days shorter), while the stage of stigma being receptive to pollen was extended by 50% compared to *C* (2 [0;5] days longer; Table S10).

Male flower anthers were visible and tended to released pollen later in *I* (although non-significantly in 2018, 2 [−3;7] days, but see Fig. S8c for other years), earlier in *IW* (−5 [−10;-0.4] days) and tended to be earlier in *W* (−4 [−10;2] days) (i.e., when 50% of C reached the pollen released stage, 27% in *I*, 94% in *IW* and 67% in *W* had reached it; Figs. 5j-k, S8c and Table S4), However, anther senescence was rather synchronous across treatments in 2018 (Fig. 5l). This suggested a double longer duration of pollen release in *IW* (5 [1;10] days longer), but a 40% shorter duration in *I* (−2 [−3;0] days shorter; Table S10).

There was evidence of a positive interaction effect between *I* and *W* for the phenophases of early leaf development, seed dispersal and anther development. The addition of warming reversed the effect of *I*, advancing phenophases even more than under *W* alone (Table S9).

## 4. Discussion

This high Arctic tundra experiment has demonstrated how icing can initially delay, but eventually enhance plant growth of a mesic community later in the growing season (Fig. 3i), while affecting reproduction (i.e., inflorescence production, Fig. 3p). This was confirmed at the species level, where the vegetative phenology of the dominant vascular plant, the dwarf shrub *S. polaris*, was delayed in early summer (Fig. 4a-b), but this seemed compensated for through an accelerated leaf development, increased SLA (Fig. 3o) and, to some extent, higher relative abundance. Seed maturation was however delayed (Fig. 4i). These observed species-and community-level delays in phenology and productivity due to icing appeared to result from later spring thawing processes, with a month-long lag in soil warming in the upper active layer (Fig. 2). The addition of summer warming partly counteracted icing effects (i.e., additive effects), but could also reverse the delayed phenology of icing by advancing it even more than under warming alone (i.e., interaction effects). This resulted in a greater increase of primary production, while also markedly advancing e.g., seed dispersal. This first multiyear tundra icing experiment also indicates a rather resistant vegetation community to a warming climate, with the absence of dramatic responses or amplified treatment effects across years.

**Fig. 4.**
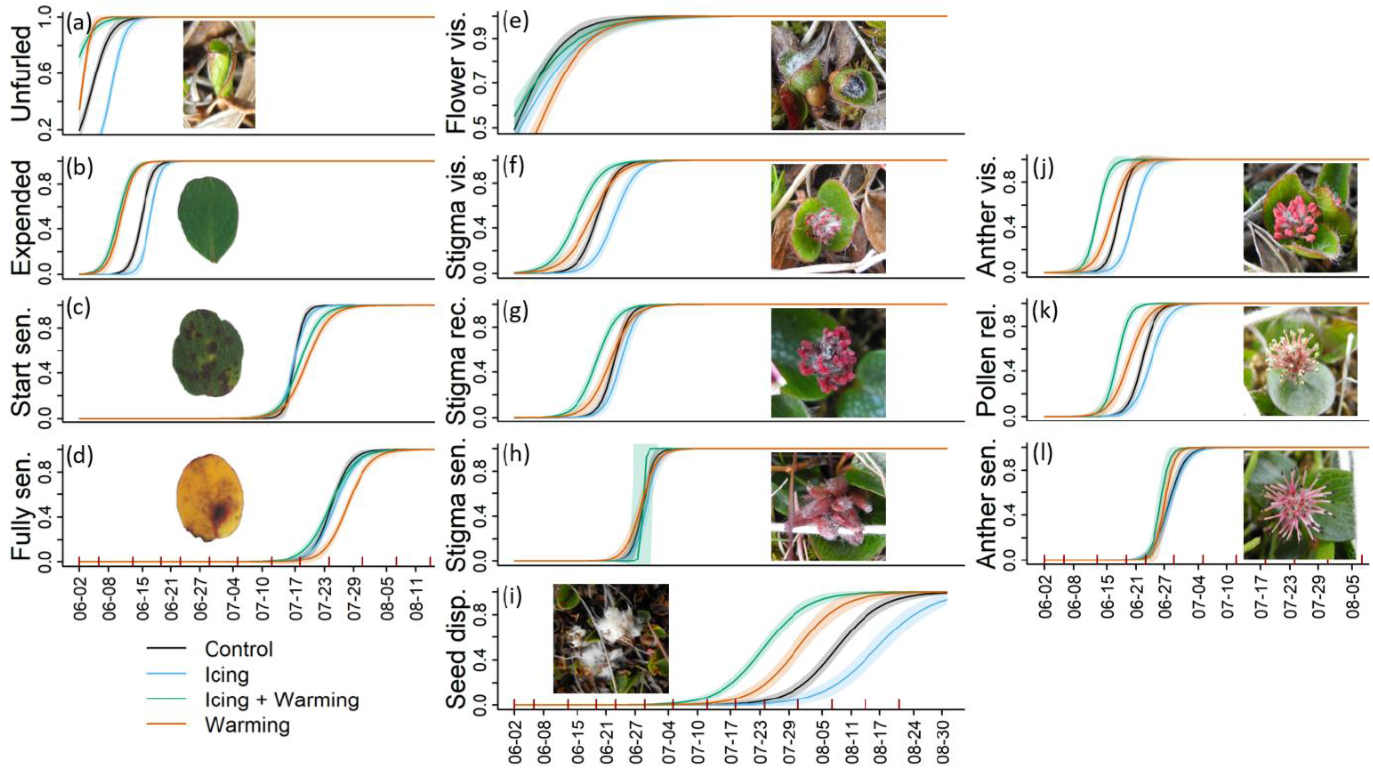
Probability curves of *Salix polaris* phenophases (i.e. the estimated probability of reaching a specific phase) for vegetative phases (a-d) and reproductive phases (e-i: female flowers; j-l: male flowers). Phenophases correspond to (a) leaves unfurled, (b) leaves fully expanded, (c) leaves started senescing, (d) leaves fully senesced, (e) flower visible (female or male), (f) stigma visible, (g) stigma receptive, (h) stigma senesced, (i) seed dispersed, (j) anther visible, (k) pollen released, and (l) anther senesced. Estimates are shown for 2018, i.e. the year when all phases could be estimated after the most intensive monitoring effort (for other years see Fig. S8). The predicted estimates were computed with generalized linear mixed-effect models, accounting for the replicated structure of the study design (i.e., random effects), and the shaded areas represent the associated 95% CIs to the fixed-effects. Red tick marks on the x-axis represent sampling days. See Fig. S9 and Table S9 for estimates of the day at which each phenophases was reached (i.e., proportion of 1).

### Delayed but increased primary production: evidence of compensatory growth?

The enhanced above-ground community-level productivity and apparent absence of major die-off due to icing contrasts with the documented ‘Arctic browning’ following extreme winter warm spells, a phenomenon related to damage and mortality in evergreen shrubs, and associated reductions in measured primary production (Bjerke et al., 2017; Frost et al., 2019; Phoenix & Bjerke, 2016). These differences may be explained by contrasting overwintering strategies of the dominant growth forms in the different studies. For instance, evergreen shrubs, virtually lacking in our study plots, maintain long-lived leaves and dormant buds well above-ground, making them more dependent on insulating snow cover and more exposed to direct damage from icing than deciduous shrubs (Givnish, 2002). Dead shoots of evergreen shrubs will remain for several years, and even if growth may be enhanced in remaining shoots (Milner et al. 2016), the net result is likely a decrease in NDVI (Bjerke et al., 2017; Treharne et al., 2020). In contrast, in our experiment dominated by the deciduous shrub *S. polaris*, there was no evident sign of dead shoots, likely allowing compensatory growth to enhance NDVI later in the growing season. Hence, despite the severity of the consecutive treatments experienced, the vegetation community studied here seemed rather resistant to the increasing frequency of icing, possibly even enhancing ‘Arctic greening’.

In the ‘ race’ for high primary production within the reduced time window available in icing plots (i.e., due to delayed soil processes), *S. polaris* appears to compensate for delayed leaves phenology by accelerating the rate of leaves development while reducing leaf dry weight and area. Because the dry weight decreased more than the area, leaf SLA (i.e., the inversed ratio) was increased over time. Leaf SLA is one of the leaf resource capture and retention traits sensitive to winter temperatures (Bjorkman et al., 2017), also used as indicator of leaf growth-rate (Pérez-Harguindeguy et al., 2013). Our findings may reveal increased growth rate from direct icing stress on the overwintering buds or due to the compressed growing season. These results agree with a laboratory icing experiment on *S. polaris* that also found reduced leaf sizes, as well as increased leaf numbers (not measured here), while photosynthetic capacity remained unchanged (Bjerke et al., 2018b). The possible increased number of leaves supports that this shrub has a remarkable ability to activate dormant buds to compensate for damages, as also shown in response to herbivory (Skarpe & Van der Wal, 2002). Furthermore, *Salix polaris* has most of its structure and reserves below-ground, accumulated in late summer as secondary growth (i.e. ring growth). Rain-on-snow events were found to reduce ring growth in coastal areas of Svalbard frequently exposed to icing (Le Moullec et al., 2020). Therefore, greater primary growth may come at the cost of reduced secondary growth, which could alter the otherwise strong correlation between primary and secondary growth in this species (Le Moullec et al., 2019).

The observed phenological delay under icing was likely caused by the extensive icing-induced lag in the warming of the upper soil layers, despite an almost simultaneous melting of ice and snow across treatments. This can be explained by infiltration and refreezing of meltwater in the underlying soil, resulting in higher ice content in near-surface soils. The thermal soil properties are strongly dependent on ice and water content. Ice-rich/wet soils require more accumulated energy to warm up than quickly thawed/dry soils, especially with presence of underlying permafrost (Isaksen et al., 2022a). Thereafter, the addition of summer warming shortened the lag in soil temperature increased found under icing.

As we expected, additive effects of summer warming (partly) counteracted negative effects of icing alone, but also with evidence of positive interaction effects. For instance, leaf phenology of *S. polaris* was instead advanced, tending to increase primary production already early in the growing season. Results from the low Arctic, where summer temperatures are much higher, match those found for our combined icing-warming treatment, i.e., simulated icing also advanced shrubs’ early leaf emergence, explained by compensatory mechanisms from frost damaged ramets (Preece et al., 2012; Preece et al., 2014). Later in the growing season, the primary production in this treatment remained high. This is likely occurring because leaves from *S. polaris* stayed fully expanded 20% longer time, and the relative abundance of the dominant graminoid *A. borealis* increased, compared to controls. In another high Arctic experiment, grasses were the growth form most rapidly responding to experimentally increased soil temperature, increasing their total live plant biomass (Brooker & van der Wal, 2003). Although there was no change in primary production under icing-warming in 2020, this can be caused by NDVI reaching saturation across treatments and controls, as expected with values approaching 0.8 (Myers-Smith et al., 2020).

The greater primary production in the second part of the growing season under icing and icing-warming combined occurred despite of a synchronous leaf senescence across treatments. This synchronicity may in part be driven by cues fixed in time, e.g., photoperiod. However, the annual variation in senescence time suggests the presence of interactions between fix cues and other variables shared across treatments, e.g., snow/ice-melt timing (Arft et al., 1999; Bjorkman et al., 2015; Cooper et al., 2011; Kelsey et al., 2021; Wipf & Rixen, 2010). A slight delay in leaf senescence under simulated warming alone was also detected in other Arctic sites, yet the main circumpolar pattern was the absence of change (Bjorkman et al., 2020; Collins et al., 2021).

### Negative effects on reproduction: a result of trade-offs?

Compensatory vegetative growth may come at the cost of reproduction, which has been observed after thaw-freeze cycles (Bokhorst et al., 2011), snow manipulation (Cooper et al., 2011), nutrient addition (Petraglia et al., 2013), and icing of evergreen shrubs (Milner et al 2016). The reduced inflorescence numbers due to icing (halved in some years) could result from prioritized allocation to vegetative growth, if not from direct winter damages. In the first experimental year, no inflorescence reduction occurred for *S. polaris*, suggesting that inflorescence buds formed the previous summer survived a winter of icing. Other species, e.g., *B. vivipara* and *A. boralis*, exhibited a strong reduction in inflorescence numbers without a lag. In an experiment conducted in the same valley, a record low number of inflorescences was associated with extreme natural rain-on-snow events in winter 2012 (Semenchuk et al., 2013). Furthermore, shorter time between green-up and flowering (i.e., green-up being more delayed than flowering) can lead to lower investment into flowering (Gougherty and Gougherty, 2018; Collins et al., 2021). Hence, similarly to other Svalbard studies, this resulted in delayed flower phenology and seed dispersal, as well as reduced flowering (Cooper et al., 2011; Karlsen et al., 2014; Semenchuk et al., 2013; Semenchuk et al., 2016).

The addition of summer warming to icing reduced the negative effects of icing on reproductive traits to a similar level or even beyond the rates obtained from warming alone. Advancing flowering and fruiting due to warming are consistent with results of simulated summer warming at a circumpolar scale (Bjorkman et al., 2020; Collins et al., 2021; Prevéy et al., 2019). Thus, like Collins et al. (2021), we found a larger shift in reproductive than vegetative phenophases, resulting in a shorter period between leaf emergence and seed dispersal. However, contrary to their overall result and to a neighbour snow manipulation experiment (Semenchuk et al., 2016), the duration of phenophases were not fixed. The duration of inflorescence and seed development were both shortened under summer warming (regardless of icing). Yet, the duration of the critical phases for reproduction, i.e., pollen release and stigmas receptive to pollen, were doubled, and this, only under combined icing-warming. Phenology studies rarely separate this short, but critical stage from the flowering time, while our results suggested important implications for pollination success (Song and Saavedra, 2018). Combining icing-warming thus indicates a high energy investment to reproduction, despite the also high investment to vegetative growth. Exploring warming effects through annual variation supported this, i.e., during the warm summer of 2020, we observed simultaneously a record high number of inflorescences and NDVI maximum.

### Evolutionary implications

Evolutionary consequences can sometimes be expected from changing trade-offs influencing sexual reproductive structure. However, the plant species composing Arctic mesic communities are generally perennial, allowing them to wait for favourable conditions to reproduce sexually (Bazzaz et al., 1987). In unfavourable years, loss of those cohorts can be buffered by seed banks, although for instance seeds from *S. polaris* are short-lived (Cooper et al., 2004) and often need to germinate within the same year they are produced. Nevertheless, *S. polaris*, as most Arctic species, also reproduces asexually by vegetative clonal growth (Jónsdóttir, 2011), and if clone size would increase, the subsequent self-pollination increase could limit evolutionary consequences of decreased seed production from environmental changes (Barrett, 2015). Furthermore, the lack of accentuated effects after five consecutive years of extreme winter conditions indicates that species in such communities have evolved strategies to cope with these conditions. Local adaptation should not be underestimated, as shown by a high Arctic transplant experiment where endogenous population were better adapted to local conditions than foreign populations, regardless of the drastic ~ 3°C increase of the summer warming treatment (Bjorkman et al., 2017).

This overall resistant and resilient mesic community studied may already have adapted to icing. In the oceanic climate of Svalbard, winter rain-on-snow events have indeed been occurring at least for decades (Vikhamar-Schuler et al., 2016). However, their frequency and extent have increased dramatically in recent years (Hansen et al., 2014; Peeters et al., 2019), and the selection acting upon life-history traits might change when acute stress episodes from occasional icy winters become ‘chronic’, as expected in the near future (Bintanja & Andry, 2017). Regions that have not yet encountered rain-on-snow events but are predicted to in the near future (e.g., Canadian high Arctic), may experience more drastic effects, especially since there would be no possible introgression of adapted genes from nearby habitat (Bjorkman et al., 2017). Comparative ‘space-for-time’ studies including Arctic regions where rain-on-snow events are still rare could shed light on plants’ adaptive capacities to icing.

### Ecological implications

The effect sizes of winter icing *versus* summer warming were of similar magnitude across most traits and within the range of natural inter-annual variation, which is typically large in the high Arctic. Although the use of open top chambers only increased summer surface air and sub-surface soil temperatures by 0.8 and 0.9 °C, respectively, this increase was similar to the decadal trend found in this region (Etzelmüller et al., 2020, Isaksen et al., 2022). It was also comparable to other ITEX sites in the high Arctic, where irradiance is low, restricting the warming effect (Bokhorst et al., 2013). There was no evidence of exacerbated treatments effects over the years, as decadal simulated summer warming also indicated (Barrett and Hollister, 2016). The observed primary production increase under warming was therefore limited. Other tundra studies/experiments that have investigated effects of altered winter conditions, notably snow accumulation, also found similar or larger effect sizes of the winter than the summer season (Cooper et al., 2011; Niittynen et al., 2020; Wahren et al., 2005). Our study thus supports the overall insight that conditions during the ‘dormant season’ can have a substantial effect on the growth of tundra vegetation.

In contrast to evergreen shrubs communities, the community studied here represent important resources for Arctic herbivores (Chapin III et al., 1996). Thus, the affected phenology and unexpected positive effect of icing on primary production may have potential implications for the whole ecosystem, such as through plant-herbivore interactions (e.g. phenological mismatches, Gauthier et al., 2013; Post & Forchhammer, 2008; Post et al., 2009), carbon fluxes (Schuur et al., 2015) and vegetation-climate feedbacks (Zhang et al., 2020). Tundra icing may thus have counteracting effects on herbivores, directly by limiting food accessibility during winter (Forbes et al., 2016; Hansen et al., 2013), but also indirectly by affecting – and possibly increasing – the food quantity during summer.

### Conclusion and outlooks

High Arctic environments are naturally variable, and plants have developed elaborate strategies to persist under these extreme conditions. Nevertheless, this study has shown that winter rain-on-snow, and associated tundra icing, can have similarly large impacts as summer warming. On the occasion of the 30-year anniversary of ITEX experiments conducted across the poles, providing critical knowledge regarding tundra responses to summer warming (Bjorkman et al., 2018; Collins et al., 2021; Prevéy et al., 2019), we call for the implementation of similar large-scale experiments investigating effects of winter warming, and rain-on-snow events in particular. Further exploration can be numerous: across communities, plant functional types, and species, as well as modulating the timing, severity, and duration of these events (Bokhorst et al., 2011; Preece & Phoenix, 2014), in interaction with different levels of summer warming, snow/ice-melt timing (Kelsey et al., 2021) and herbivory (Gornall et al., 2009). Such a coordinated effort on long-term effects is required to gain a holistic understanding of future changes in the tundra biome and its possible resistance and resilience to both winter and summer warming (Bjorkman et al., 2020). This is especially urgent given that winter warming is by far outpacing summer warming (Isaksen et al., 2022b), with potentially ecosystem-wide consequences such as alteration of below-ground vegetation productivity, microorganism activity, and carbon flux exchange, and cascading effects across the food web. This pioneer long-term icing experiment sets the stage for forthcoming studies to fill this fundamental knowledge gap.

## Acknowledgements

This project was funded by the Research Council Norway (FRIPRO 276080 and SFF-III 223257) and Svalbard Environmental Protection Fund (project 16/113). We thank UNIS logistics for logistical support and Longyearbyen Lokalstyre for permission to perform the experiment (RiS ID: 10484). We are also deeply grateful for all the help in the field from Solvei B. Hovdal, Martha Grotheim, Hanne K. Haraldsen, Julia Greulich, Karrín Björnsdóttir, Kristine Valøen, Robin Zweigel, Ådne Nafstad, Marit Arneberg, Kristiane Midhaug, Marte Søreng, Marianne Angård, Hamish Burnett, Lia Lechler, Charlotte Karlsen, Kjerstin Hilmarsen, Lukas Tietgen, and Svea Zimmermann.

## Notes

### Competing Interest Statement

The authors have declared no competing interest.

### Summary of Updates

Revised version of the text and illustrations. The supplementary files are now uploaded.

