## Supplementary material for "Towards rainy high Arctic winters: how ice-encasement impacts tundra plant phenology, productivity and reproduction"

### Supplementary material to: Towards rainy Arctic winters: experimental icing impacts tundra plant productivity and reproduction

Mathilde Le Moullec, Anna-Lena Hendel, Matteo Petit Bon, Ingibjörg Svala Jónsdóttir, Øystein Varpe, René van der Wal, Larissa Teresa Beumer, Kate Layton-Matthews, Ketil Isaksen and Brage Bremset Hansen.

<u>Table of content:</u>

Table S1. Summary statistics of summer soil temperature and volumetric water content.

Table S2. Sampling dates and number of repeated measurements.

Table S3. Correlation coefficients between maximum NDVI and relative abundance.

Table S4. Summary statistic of NDVI metrics.

Table S5. Summary statistics of relative abundance.

Table S6. Summary statistics of leaf traits.

Table S7. Summary statistics of flower counts.

Table S8. Summary statistics of measurements' temporal trends.

Table S9. Summary statistic of the day-of-year a certain phenophase is reached.

Table S10. Summary statistic of time spent in a certain phenophase

Figure S1. Mesic habitat species composition.

Figure S2: Immediate icing treatment effect on soil temperature.

Figure S3. Estimated NDVI curves across treatments and years.

Figure S4. Annual relative abundance.

Figure S5. Ordination of the mesic community species composition.

Figure S6. Leaf size traits of *Salis polaris* across years and treatments.

Figure S7. Flower counts per year, treatment and species/group.

Figure S8. Salix *polaris* phenology.

Figure S8. Phenophases' density plots.

Table S1. Summary statistics of summer air and soil temperature and volumetric water content (i.e., soil moisture in %). Surface air temperature (5 cm high) was recorded with HOBO loggers (type U23-003/UA-001; ± 0.2°C accuracy) every 30 min in 2016-2018 and averaged daily over mid-June to end-August before further modelling. This comprised two plots in 2016-2017 and 6 plots in 2018. Plot-level soil sub-surface temperature (at 2 and 5 cm depth in each 36 plots) was recorded with iButton loggers (type DS1921G-F5, ± 1.0 °C accuracy) every 240 minutes and averaged daily over June-August before further modeling. Soil moisture was measured with a ML3 ThetaProbe Sensor (HH2 Soil Moisture Meter from Delta-T Devices Ltd, UK, 1% accuracy, 5-10 cm depth) via repeated measurements at five points per plot, 3-15 times per summer (rounds). a) Summary table with soil temperature and moisture model prediction and 95% CI per treatment and year. Model 1 included treatment [category] × year [factor] interaction as fixed effects. Model 2 included only treatment, while year was set a random intercept (2016, 2018 and 2019 for temperature and 2016-2020 for moisture), with date nested within year. From these models, we documented the predicted means for controls [*C*] and effect sizes for treatments, in contrast to *C*. We reported the random intercept effects corresponding to Model 2. b) Pearson correlation coefficients between soil moisture and sub-surface (5 cm depth) temperature for years with repeated measurements across the growing season (2018-2019, df = 18). To compute these correlation coefficients we first fitted separate linear mixed-effect models for moisture and temperature, with treatment × day-of-year [factor] included as fixed effect. We used temperatures recorded on the same days as moisture measurements were conducted.  The random intercept structure accounted for plots nested within blocks. Then, the extracted fixed-effect estimates for each variable and correlated them with each other. I = icing, IW = icing × warming, W = warming.

a)

| *Model* | *Treatment* | *Year* | *T°C (5 cm high)* | *T°C (2 cm depth)* | *T°C (5 cm depth)* | *Moisture %* |
|---|---|---|---|---|---|---|
| 1 | C | 2016 | 7.7 [7.11,8.22] | 6 [5.5,6.6] | 5.1 [-0.5,10.8] | 32.8 [26.7,39] |
| | I | 2016 | - | 5.9 [5.3,6.5] | 5.1 [-0.6,10.7] | 36.6 [30.5,42.7] |
| | IW | 2016 | - | 7.2 [6.6,7.8] | 5.5 [-0.1,11.2] | 31.9 [25.8,38] |
| | W | 2016 | 8.4 [7.9,9.0] | 6.9 [6.4,7.5] | 5.4 [-0.2,11] | 33.8 [27.7,40] |
| | C | 2017 | 6.9 [6.5,7.4] | - | - | 32.6 [27.1,38.1] |
| | I | 2017 | - | - | - | 32.8 [27.3,38.3] |
| | IW | 2017 | - | - | - | 30.6 [25.1,36.1] |

| | | | | | | |
|---|---|---|---|---|---|---|
| | W | 2017 | 7.7 [7.2,8.1] | - | - | 34.2 [28.8,39.7] |
| | C | 2018 | 7.3 [6.8,7.7] | 6 [5.4,6.6] | 5 [-0.6,10.6] | 33 [27.5,38.4] |
| | I | 2018 | - | 5.9 [5.3,6.5] | 5 [-0.6,10.7] | 32.6 [27.1,38] |
| | IW | 2018 | - | 6.7 [6.1,7.2] | 5.6 [0,11.3] | 28.5 [23.1,34] |
| | W | 2018 | 8.0 [7.6,8.5] | 6.9 [6.4,7.5] | 5.9 [0.2,11.5] | 32 [26.5,37.4] |
| | C | 2019 | - | 7.2 [6.6,7.8] | 5.4 [-0.3,11] | 27.3 [21.9,32.8] |
| | I | 2019 | - | 6.7 [6.1,7.3] | 4.9 [-0.8,10.5] | 27.9 [22.4,33.4] |
| | IW | 2019 | - | 7.4 [6.8,8] | 5.5 [-0.1,11.2] | 23.9 [18.4,29.3] |
| | W | 2019 | - | 8.4 [7.8,9] | 6 [0.4,11.7] | 27.5 [22,33] |
| | C | 2020 | - | - | - | 36.5 [31,42.1] |
| | I | 2020 | - | - | - | 36 [30.4,41.5] |
| | IW | 2020 | - | - | - | 33 [27.5,38.6] |
| | W | 2020 | - | - | - | 35.7 [30.1,41.2] |
| 2 | C | 16-20 | 7.3 [6.4,8.2] | 6.4 [5.7,7.1] | 4.6 [3.7,5.6] | 34.6 [28.9,40.2] |
| 2 | I - C | 16-20 | - | -0.2 [-0.4,0.1] | -0.1 [-0.4,0.2] | 0 [-3.6,3.6] |
| | IW - C | 16-20 | - | 0.8 [0.6,1.1] | 0.4 [0.1,0.7] | -3.6 [-7.2,0.1] |
| | W - C | 16-20 | 0.8 [0.7,0.9] | 0.9 [0.7,1.2] | 0.6 [0.3,1] | -0.1 [-3.7,3.5] |
| 2 | Residuals | | 0.23 | 0.49 | 0.32 | 47.11 |
| | blocks/plots/sub-squares | | - | - | - | 19.47 |
| | blocks/plots | | - | 0.12 | 0.20 | 17.76 |
| | block | | - | <0.01 | 0.03 | 9.16 |
| | year/day of year | | 4.29 | 4.53 | 3.75 | 37.21 |
| | year | | 0.08 | 0.29 | <0.01 | 7.06 |
| | Observations | | 406 [a] | 7031[b] | 7353[b] | 5808[c] |

[a] Monitoring for 3 summer each day (15th June-1st Sep, n = 68) in two plots.
[b] 36 observations (12 plots within 3 blocks) each day (n = 93), over 3 summers.
[c] 180 observations (5 sub-squares, within 12 plots, within 3 blocks) each round (n = 35) distributed over 5 years. The first year (2016) had no sub-square replication.

b)

| Treatment | Correlation coefficient [95% CI] |
|---|---|
| C | -0.69 [-0.87:-0.35] |
| I | -0.79 [-0.92:-0.49] |
| IW | -0.71 [-0.88:0.40] |
| W | -0.60 [0.82:-0.21] |

Table S2. Sampling dates and number of measurement repeats (n). Abundance measurements represent the relative abundance measured by the point intercept method. For the *Salix polaris* leaf collection, the protocols varied between years: 'Random' indicates that *S. polaris* leaves were gathered from five shoots collected at random (i.e., at five sub-square intercepts of the frame) and in 2018, the leaf's respective shoot ID was available and used in the linear mixed-effect modelling; 'Largest' indicates that the biggest *S. polaris* leaf was collected in each 16 sub-squares per plot.

| | 2016 | 2017 | 2018 | 2019 | 2020 |
|---|---|---|---|---|---|
| Date of icing | 3 – 4 Feb | 22 – 23 Jan | 4 – 5 Jan | 24 – 25 Feb | 18 – 19 Feb |
| Date OTC setup | 23 May | 31 May | 20 May | 4 Jun | 20 May |
| NDVI rounds | 23 Jun – 11 Aug (n = 9) | 21 Jun – 30 Jul (n = 7) * | 01 Jun – 20 Aug (n = 15) | 11 Jun – 17 Aug (n = 9) | 10 Jun – 18 Aug (n = 10) |
| Phenology rounds | 23 Jun – 13 Aug (n = 12) | 22 Jun – 31 Jul (n = 7) | 02 Jun – 21 Aug (n = 14) | 07 Jun – 10 Aug (n = 5) | NA |
| Flower counts | 14 Jul | 25 July | 17 Jul | 16 Jul | 9 Jul |
| Relative abundance | 2 – 4 Aug | 4 – 6 Aug | 30 – 31 Jul | 23 – 24 Jul | NA |
| *S. polaris* leaf collection | 5 Aug | 2 Aug | 30 Jul | 30 Jul | 28 Jul |
| *S. polaris* leaf collection protocol | Random | Random | Random | Largest | Largest |

* too short summer period to be included in the analysis

Table S3. Pearson's correlation coefficients between plot-specific maximum NDVI and relative abundance estimates (assessed via point intercept method), separately for the two dominant vascular plant species (*Salix polaris, Alopecurus borealis*), different groups of vascular plant species, and bryophytes.

| | 2016 (df = 34) | 2018 (df = 34) | 2019 (df = 34) | Total (df = 106) |
|---|---|---|---|---|
| *S. polaris* | 0.45 [0.14,0.68] | 0.30 [-0.03,0.57] | 0.46 [0.16,0.69] | 0.40 [0.23,0.55] |
| *A. borealis* | 0.32 [-0.01,0.58] | 0.23 [-0.11,0.52] | -0.10 [0.42,0.24] | 0.15 [-0.04,0.33] |
| Vascular plants | 0.42 [0.11,0.66] | 0.38 [0.06,0.63] | 0.36 [0.04,0.62] | 0.36 [0.19,0.52] |
| Vascular plants without *S. polaris* and *A. borealis* | 0.15 [-0.19,0.46] | 0.07 [-0.27,0.39] | 0.04 [-0.29,0.36] | 0.09 [-0.10,0.28] |
| Bryophytes | -0.40 [-0.64,-0.08] | -0.22 [-0.51,0.11] | -0.00 [-0.33,0.33] | -0.19 [-0.37:-0.00] |

Table S4. Summary statistic of NDVI metrics presented in Figure 3a-e and 3i-m (see main text). Model predictions were computed for controls [*C*], which are the reference level for the treatments' effect sizes reported, for each year category respectively. Model predictions and estimates are presented with their associated 95% confidence intervals. Different models were fitted to quantify within (model 1) or across (model 3 and 5) years effects. Model 1 included 'treatment' (as categories; *C*, *I* = icing, *W* = warming, *IW* = icing and warming combined) in interaction with 'year' (as factor) as fixed effects (treatment × year). Model 3 and 5 included only 'treatment' (as categories) as fixed effects while 'year' was set as random intercept, including all years with NDVI measurements in model 3 (2016, 2018, 2019, 2020) and common years with all types of measurements in model 5 (2016, 2018, 2019). The random intercept effects reported are associated to model 3. Model 2, 4 and 6 tested for the interaction between *I* and *W* treatments (binomial variables, *I* × *W*). In all models, the random intercept structure included 12 plots, nested within 3 blocks, over 4 years (= 144 observations, except for models 5 and 6 with 3 years). Estimates in bold have CIs non-overlapping with 0. Time of maximum NDVI (Time of max.) is expressed in day-of-year unit, maximum NDVI (max.) and the cumulative NDVI metrics (cum.) can take values from (-1 to 1).

| *Model* | *Treatment/ Contrast* | *Year* | *Time of max.* | *Max.* | *Cum. start* | *Cum. end* | *Cum. tot* |
|---|---|---|---|---|---|---|---|
| 1a | C | 2016 | 197 [189,205] | 0.742 [0.7154,0.7685] | 0.71 [0.69,0.73] | 0.71 [0.69,0.73] | 0.71 [0.69,0.73] |
| 1b | I - C | 2016 | 4 [-1,9] | 0.0298 [-0.0023,0.0619] | 0.01 [-0.02,0.04] | **0.04 [0.01,0.07]** | **0.03 [0,0.05]** |
| | IW - C | 2016 | -1 [-5,4] | 0.0351 [0.003,0.0673] | **0.04 [0.01,0.06]** | **0.05 [0.02,0.08]** | **0.04 [0.02,0.07]** |
| | W - C | 2016 | -3 [-8,1] | -0.0028 [-0.0349,0.0294] | 0.01 [-0.02,0.04] | 0.01 [-0.02,0.04] | 0.01 [-0.02,0.04] |
| 2 | I x W | 2016 | -1 [-8,6] | 0.0081 [-0.0373,0.0535] | 0.02 [-0.03,0.06] | 0 [-0.04,0.04] | 0.01 [-0.03,0.05] |
| 1a | C | 2018 | 199 [191,207] | 0.7556 [0.729,0.7821] | 0.65 [0.63,0.67] | 0.71 [0.69,0.73] | 0.67 [0.66,0.69] |
| 1b | I - C | 2018 | 4 [-1,9] | 0.0143 [-0.0179,0.0464] | 0 [-0.03,0.03] | **0.03 [0,0.06]** | 0.01 [-0.02,0.04] |
| | IW - C | 2018 | -1 [-6,4] | 0.018 [-0.0141,0.0502] | **0.03 [0.01,0.06]** | 0.03 [-0.01,0.06] | **0.03 [0,0.05]** |
| | W - C | 2018 | -4 [-9,1] | -0.0048 [-0.0369,0.0273] | 0.01 [-0.02,0.04] | -0.01 [-0.04,0.03] | 0.01 [-0.02,0.03] |
| 2 | I x W | 2018 | -1 [-8,6] | 0.0086 [-0.0369,0.054] | 0.02 [-0.02,0.06] | 0 [-0.04,0.05] | 0.01 [-0.03,0.05] |

| | | | | | | | |
|---|---|---|---|---|---|---|---|
| 1a | C | 2019 | 192 [184,200] | 0.7571 [0.7305,0.7836] | 0.65 [0.63,0.67] | 0.71 [0.69,0.74] | 0.69 [0.67,0.7] |
| 1b | I - C | 2019 | **6 [1,11]** | 0.0036 [-0.0285,0.0357] | -0.01 [-0.04,0.02] | 0.02 [-0.01,0.05] | 0.01 [-0.02,0.04] |
| | IW - C | 2019 | **7 [2,12]** | 0.0126 [-0.0195,0.0447] | <0.01 [-0.03,0.03] | **0.03 [0,0.06]** | 0.02 [-0.01,0.04] |
| | W - C | 2019 | 3 [-2,8] | -0.0004 [-0.0325,0.0317] | <0.01 [-0.03,0.03] | <0.01 [-0.03,0.04] | <0.01 [-0.02,0.03] |
| 2 | I x W | 2019 | -2 [-9,5] | 0.0094 [-0.036,0.0548] | 0.01 [-0.03,0.05] | <0.01 [-0.04,0.04] | <0.01 [-0.03,0.04] |
| 1a | C | 2020 | 196 [188,204] | 0.785 [0.7585,0.8116] | 0.68 [0.66,0.7] | 0.74 [0.72,0.76] | 0.71 [0.69,0.73] |
| 1b | I - C | 2020 | 3 [-1,8] | -0.0096 [-0.0418,0.0225] | -0.02 [-0.05,0.01] | <0.01 [-0.03,0.03] | -0.01 [-0.04,0.02] |
| | IW - C | 2020 | 0 [-4,5] | -0.0127 [-0.0449,0.0194] | -0.02 [-0.05,0.01] | -0.01 [-0.04,0.02] | -0.01 [-0.04,0.02] |
| | W - C | 2020 | -6 [-11,-1] | -0.0058 [-0.0379,0.0264] | 0.02 [-0.01,0.05] | 0 [-0.03,0.03] | 0.01 [-0.02,0.04] |
| 2 | I x W | 2020 | 3 [-4,10] | 0.0026 [-0.0428,0.0481] | -0.02 [-0.06,0.02] | -0.01 [-0.05,0.03] | -0.01 [-0.05,0.02] |
| 3a | C | 16-20 | 196 [187,205] | 0.7599 [0.7336,0.7862] | 0.67 [0.62,0.72] | 0.72 [0.7,0.74] | 0.7 [0.66,0.73] |
| 3b | I - C | 16-20 | **4 [2,7]** | 0.0095 [-0.0195,0.0385] | -0.01 [-0.03,0.02] | **0.02 [0.01,0.05]** | 0.01 [-0.02,0.03] |
| | IW - C | 16-20 | 2 [-1,4] | 0.0133 [-0.0157,0.0422] | 0.01 [-0.01,0.04] | **0.02 [0.01,0.05]** | 0.02 [-0.01,0.04] |
| | W - C | 16-20 | -3 [-5,0] | -0.0034 [-0.0324,0.0255] | 0.01 [-0.01,0.04] | <0.01 [-0.03,0.03] | 0.01 [-0.02,0.03] |
| 4 | I x W | 16-20 | 0 [-4,3] | 0.0072 [-0.0338,0.0482] | 0.01 [-0.03,0.04] | <0.01 [-0.04,0.04] | <0.01 [-0.03,0.04] |
| 5a | C | 16-18-19 | 196 [188,204] | 0.7515 [0.7286,0.7745] | 0.67 [0.58,0.76] | 0.71 [0.69,0.73] | 0.69 [0.64,0.74] |
| 5b | I - C | 16-18-19 | **5 [1,8]** | 0.0159 [-0.0125,0.0443] | <0.01 [-0.02,0.02] | **0.03 [0,0.06]** | 0.01 [-0.01,0.04] |
| | IW - C | 16-18-19 | 2 [-1,5] | 0.0219 [-0.0064,0.0503] | **0.02 [0,0.04]** | **0.03 [0.01,0.06]** | **0.03 [0,0.05]** |
| | W - C | 16-18-19 | -2 [-5,2] | -0.0026 [-0.031,0.0257] | 0.01 [-0.02,0.03] | <0.01 [-0.03,0.03] | 0.01 [-0.02,0.03] |
| 6 | I x W | 16-18-19 | -1 [-6,3] | 0.0087 [-0.0314,0.0488] | 0.01 [-0.02,0.05] | <0.01 [-0.04,0.04] | 0.01 [-0.03,0.04] |
| 3 | Residuals | | 27.73 | 0.0004 | 0.0005 | 0.0004 | 0.0003 |

| | | | | | |
|---|---|---|---|---|---|
| blocks/plots | 0.60 | 0.0008 | 0.0005 | 0.0008 | 0.0006 |
| blocks | 17.36 | 0.0001 | <0.0001 | <0.0001 | <0.0001 |
| year | 2.28 | 0.0001 | 0.0012 | <0.0001 | 0.0004 |
| Observations | 144 | 144 | 144 | 144 | 144 |

Table S5. Summary statistic of relative abundance presented in Figure 3f, 3n (see main text) and S3. Relative abundance corresponds to the number of hits per plots measured with the point intercept method for all vascular plants of the mesic community and separately for the two most abundant species, *Alopecurus borealis* and *Salix polaris*. Model predictions were computed for controls [*C*], which are the reference level for the treatments' effect sizes reported, for each year category respectively. Different models were fitted to quantify within (model 1) or across (model 3 and 5) years effects. Model 1 included 'treatment' (as categories; *C*, *I* = icing, *W* = warming, *IW* = icing and warming combined) in interaction with 'year' (as factor) as fixed effects (treatment × year). Model 3 and 5 included only 'treatment' (as categories) as fixed effects while 'year' was set as random intercept, including all years with NDVI measurements in model 3 (2016, 2017, 2018, 2019) and common years with all types of measurements in model 5 (2016, 2018, 2019). The random intercept effects reported are associated to model 3. Model 2, 4 and 6 tested for the interaction between *I* and *W* treatments (binomial variables, *I* × *W*). In all models, the random intercept structure included 12 plots, nested within 3 blocks, over 4 years (= 144 observations, except for models 5 and 6 with 3 years). Model predictions and estimates are presented on the log-scale with their associated 95% Cis, in bold when estimates' CIs non-overlap with 0.

| *Model* | *Treatment/ Contrast* | *Year* | *Vascular plants* | *A. borealis* | *S. polaris* |
|---|---|---|---|---|---|
| 1a) | C | 2016 | 3.14 [2.95,3.32] | 1.65 [1.22,2.09] | 2.23 [1.82,2.65] |
| 1b) | I - C | 2016 | 0.16 [-0.1,0.42] | 0.25 [-0.35,0.86] | 0.17 [-0.36,0.71] |
| | IW - C | 2016 | **0.52 [0.27,0.77]** | **0.96 [0.38,1.54]** | 0.29 [-0.24,0.81] |
| | W - C | 2016 | 0.23 [-0.03,0.49] | 0.46 [-0.14,1.05] | -0.36 [-0.92,0.2] |
| 2) | I x W | 2016 | 0.14 [-0.22,0.49] | 0.25 [-0.57,1.07] | 0.47 [-0.29,1.24] |
| 1a) | C | 2017 | 3.45 [3.28,3.63] | 1.88 [1.46,2.3] | 2.39 [1.99,2.8] |
| 1b) | I - C | 2017 | -0.04 [-0.28,0.21] | **-0.82 [-1.47,-0.17]** | 0.31 [-0.21,0.83] |
| | IW - C | 2017 | 0.13 [-0.11,0.37] | 0.06 [-0.53,0.64] | 0.14 [-0.38,0.67] |
| | W - C | 2017 | 0.03 [-0.21,0.28] | -0.23 [-0.83,0.38] | -0.52 [-1.07,0.03] |
| 2) | I x W | 2017 | 0.14 [-0.21,0.48] | **1.1 [0.21,1.98]** | 0.35 [-0.4,1.1] |
| 1a) | C | 2018 | 3.42 [3.24,3.6] | 2.11 [1.7,2.51] | 2.29 [1.88,2.7] |
| 1b) | I - C | 2018 | -0.07 [-0.32,0.18] | -0.34 [-0.93,0.25] | 0.16 [-0.37,0.69] |
| | IW - C | 2018 | 0.08 [-0.16,0.33] | 0.19 [-0.38,0.75] | 0.2 [-0.32,0.73] |

| | | | | | |
|---|---|---|---|---|---|
| | W - C | 2018 | 0.07 [-0.18,0.32] | 0 [-0.57,0.57] | -0.3 [-0.85,0.25] |
| 2) | I x W | 2018 | 0.09 [-0.26,0.44] | 0.53 [-0.29,1.34] | 0.34 [-0.41,1.1] |
| 1a) | C | 2019 | 3.34 [3.16,3.52] | 1.49 [1.04,1.94] | 2.42 [2.02,2.83] |
| 1b) | I - C | 2019 | 0.08 [-0.17,0.33] | 0.1 [-0.53,0.73] | 0.09 [-0.43,0.61] |
| | IW - C | 2019 | 0.2 [-0.04,0.45] | 0.4 [-0.21,1.02] | 0.11 [-0.42,0.63] |
| | W - C | 2019 | 0.19 [-0.06,0.44] | 0.55 [-0.06,1.16] | -0.39 [-0.93,0.15] |
| 2) | I x W | 2019 | -0.07 [-0.42,0.28] | -0.25 [-1.11,0.61] | 0.4 [-0.35,1.15] |
| 3a) | C | 16-19 | 3.34 [3.19,3.49] | 1.79 [1.37,2.21] | 2.34 [1.96,2.71] |
| 3b) | I - C | 16-20 | 0.03 [-0.18,0.23] | -0.18 [-0.71,0.34] | 0.19 [-0.28,0.66] |
| | IW - C | 16-20 | **0.23 [0.03,0.43]** | 0.42 [-0.1,0.93] | 0.18 [-0.29,0.65] |
| | W - C | 16-20 | 0.12 [-0.08,0.33] | 0.18 [-0.33,0.7] | -0.39 [-0.88,0.09] |
| 4) | I x W | 16-20 | 0.08 [-0.21,0.36] | 0.41 [-0.32,1.15] | 0.39 [-0.29,1.06] |
| 5a) | C | 16-18-19 | 3.3 [3.14,3.46] | 1.76 [1.34,2.18] | 2.32 [1.97,2.66] |
| 5b) | I - C | 16-18-19 | 0.05 [-0.17,0.27] | -0.01 [-0.55,0.52] | 0.14 [-0.33,0.61] |
| | IW - C | 16-18-19 | **0.27 [0.05,0.49]** | **0.55 [0.02,1.07]** | 0.2 [-0.27,0.66] |
| | W - C | 16-18-19 | 0.16 [-0.06,0.38] | 0.31 [-0.22,0.84] | -0.32 [-0.8,0.17] |
| 6) | I x W | 16-18-19 | 0.05 [-0.26,0.36] | 0.25 [-0.49,1] | 0.37 [-0.3,1.04] |
| 3) | Residuals | | 0.0316 | 0.1400 | 0.0927 |
| | blocks/plots | | 0.0396 | 0.2749 | 0.2387 |
| | blocks | | 0.0005 | <0.0001 | 0.0197 |
| | year | | 0.0009 | 0.0392 | 0.0011 |
| | Observations | | 144 | 144 | 144 |

Table S6. Summary statistic of *Salix polaris*'s leaf traits. Specific leaf area (SLA, presented in Figure 3g and 3o) is the ratio between leaf area (Area) and its dry weight (presented in Figure S5). Single leaf area ($mm^2$) and dry weight (mg) were first log-transformed and then standardized within each year (mean = 0, sd = 1) to enable treatment effect size comparison between year. This was necessary due to the change in sampling design between years, possibly affecting the annual mean and variance. Model predictions were computed for controls [*C*], which are the reference level for the treatments' effect sizes reported, for each year category respectively. Different models were fitted to quantify within (model 1) or across (model 3 and 5) years effects. Model 1 included 'treatment' (as categories; *C*, *I* = icing, *W* = warming, *IW* = icing and warming combined) in interaction with 'year' (as factor) as fixed effects (i.e., treatment × year). Model 3 and 5 included only 'treatment' (as categories) as fixed effects while 'year' was set as random intercept, including all years with leaf traits measurements in model 3 (2016-2020) and years with all types of measurements in model 5 (2016, 2018 and 2019). The random intercept effects reported are associated to model 3. Model 2, 4 and 6 tested for the interaction between *I* and *W* treatments (binomial variables, *I* × *W*). Model predictions and estimates are presented on the log-scale with their associated 95% CIs, in bold when estimates' CIs non-overlap with 0.

| *Model* | *contrast* | *year* | *SLA* | *Area* | *Dry weight* |
|---|---|---|---|---|---|
| 1[a] | C | 2016 | 0.12 [-0.19,0.43] | 0.06 [-0.27,0.38] | 0.01 [-0.33,0.35] |
| 1[b] | I - C | 2016 | 0.12 [-0.2,0.45] | -0.03 [-0.4,0.35] | -0.06 [-0.43,0.3] |
| | IW - C | 2016 | **-0.47 [-0.78,-0.15]** | -0.21 [-0.57,0.15] | -0.01 [-0.36,0.34] |
| | W - C | 2016 | -0.02 [-0.35,0.3] | -0.11 [-0.48,0.26] | -0.09 [-0.45,0.27] |
| 2[b] | I x W | 2016 | **-0.57 [-1.09,-0.05]** | -0.07 [-0.66,0.51] | 0.14 [-0.43,0.71] |
| 1[a] | C | 2017 | -0.18 [-0.48,0.11] | 0.03 [-0.28,0.35] | 0.08 [-0.24,0.41] |
| 1[b] | I - C | 2017 | 0.11 [-0.19,0.41] | 0.08 [-0.27,0.44] | 0.03 [-0.31,0.37] |
| | IW - C | 2017 | **0.33 [0.04,0.62]** | -0.17 [-0.51,0.17] | -0.24 [-0.57,0.09] |
| | W - C | 2017 | 0.25 [-0.05,0.55] | -0.05 [-0.4,0.29] | -0.12 [-0.46,0.21] |
| 2[b] | I x W | 2017 | -0.03 [-0.51,0.45] | -0.2 [-0.75,0.35] | -0.15 [-0.68,0.38] |
| 1[a] | C | 2018 | -0.06 [-0.37,0.25] | 0.09 [-0.23,0.42] | 0.1 [-0.24,0.44] |
| 1[b] | I - C | 2018 | 0.31 [-0.01,0.63] | -0.31 [-0.67,0.06] | -0.34 [-0.7,0.01] |
| | IW - C | 2018 | -0.07 [-0.39,0.24] | -0.21 [-0.57,0.15] | -0.15 [-0.5,0.2] |
| | W - C | 2018 | -0.01 [-0.34,0.31] | 0.16 [-0.21,0.53] | 0.12 [-0.23,0.48] |

| | | | | | |
|---|---|---|---|---|---|
| 2 [b] | I x W | 2018 | -0.37 [-0.89,0.15] | -0.06 [-0.65,0.52] | 0.07 [-0.5,0.64] |
| 1[a] | C | 2019 | -0.15 [-0.46,0.16] | 0.25 [-0.08,0.58] | 0.27 [-0.07,0.6] |
| 1[b] | I - C | 2019 | **0.46 [0.15,0.78]** | **-0.45 [-0.82,-0.08]** | **-0.54 [-0.9,-0.19]** |
| | IW - C | 2019 | -0.11 [-0.42,0.21] | -0.35 [-0.71,0.01] | -0.26 [-0.61,0.09] |
| | W - C | 2019 | **0.31 [0,0.62]** | -0.34 [-0.7,0.01] | **-0.4 [-0.75,-0.06]** |
| 2 [b] | I x W | 2019 | **-0.88 [-1.39,-0.38]** | 0.44 [-0.12,1.01] | **0.69 [0.13,1.24]** |
| 1[a] | C | 2020 | -0.26 [-0.58,0.06] | 0.15 [-0.19,0.49] | 0.21 [-0.14,0.56] |
| 1[b] | I - C | 2020 | **0.52 [0.2,0.85]** | -0.24 [-0.62,0.13] | **-0.38 [-0.75,-0.01]** |
| | IW - C | 2020 | 0.04 [-0.28,0.36] | -0.06 [-0.43,0.3] | -0.06 [-0.41,0.29] |
| | W - C | 2020 | **0.54 [0.21,0.87]** | **-0.4 [-0.77,-0.03]** | **-0.52 [-0.88,-0.16]** |
| 2 [b] | I x W | 2020 | **-1.03 [-1.55,-0.5]** | **0.58 [0,1.16]** | **0.84 [0.27,1.41]** |
| 3 [a] | C | 16-20 | -0.11 [-0.38,0.16] | 0.11 [-0.19,0.4] | 0.13 [-0.18,0.43] |
| 3[b] | I - C | 16-20 | **0.28 [0,0.55]** | -0.16 [-0.49,0.17] | -0.23 [-0.55,0.09] |
| | IW - C | 16-20 | -0.02 [-0.29,0.25] | -0.19 [-0.5,0.13] | -0.15 [-0.45,0.16] |
| | W - C | 16-20 | 0.23 [-0.04,0.5] | -0.16 [-0.48,0.16] | -0.21 [-0.52,0.09] |
| 4 [b] | I x W | 16-20 | **-0.53 [-0.99,-0.08]** | 0.13 [-0.37,0.63] | 0.29 [-0.19,0.78] |
| 5 [a] | C | 16-18-19 | -0.03 [-0.3,0.23] | 0.15 [-0.17,0.46] | 0.14 [-0.18,0.45] |
| 5 [b] | I - C | 16-18-19 | **0.31 [0,0.61]** | -0.28 [-0.64,0.09] | -0.33 [-0.69,0.02] |
| | IW - C | 16-18-19 | -0.21 [-0.51,0.09] | -0.24 [-0.58,0.11] | -0.13 [-0.47,0.21] |
| | W - C | 16-18-19 | 0.06 [-0.25,0.36] | -0.07 [-0.42,0.27] | -0.09 [-0.44,0.25] |
| 6 [b] | I x W | 16-18-19 | **-0.57 [-1.06,-0.09]** | 0.11 [-0.44,0.66] | 0.29 [-0.25,0.84] |
| 3 | Residuals | | 0.88 | 0.85 | 0.85 |
| | blocks/plots | | 0.12 | 0.17 | 0.16 |
| | blocks | | 0.02 | 0.01 | 0.02 |
| | Observations | | 3002[c] | 3002[c] | 3002[c] |

[a] Model prediction
[b] Model estimate
[c] 36 plots (nested within 3 blocks) monitored over 5 years (mean standardized, random variation <0.001), with on average 16.7 leaves collected per plot (range: 11 to 45 leaves per plot).

Table S7. Summary statistic of inflorescence counts. The total counts gather the inflorescence number of *S. polaris* (female catkins and male flowers), *B. vivipara* (combining reproductive shoots with inflorescence and/or bulbils) and the ‘graminoids’ combining *A. borealis, L. confusa* and *P. arctica*. Model predictions were computed for controls [*C*], which were the reference level for the treatments’ effect sizes reported, for each year category respectively. Different models were fitted to quantify within (model 1) or across (model 3 and 5) years effects. Model 1 included ‘treatment’ [category] in interaction with ‘year’ [factor] and ‘species’ [category] as fixed effects (i.e., treatment × year × species, species was not included in the column ‘Total’). Model 3 and 5 included only ‘treatment’ (as categories) as fixed effects while ‘year’ was set as random intercept, including all years of the study in model 3 (2016-2020) and years with all types of measurements in model 5 (2016, 2018 and 2019). Model 2, 4 and 6 tested for the interaction between *I* and *W* treatments (binomial variables, *I* × *W*). Model predictions and estimates are presented on the square root-scale with their associated 95% CIs. In bold, estimates’ CIs non-overlap with 0. Backtranformed predictions and estimates are plotting in Figure 3h and 3p (total counts), as well as in Figure S6 (species/group count). *I* = icing, *W* = warming, *IW* = icing and warming combined

| | | | *Inflorescence count* | | | |
|---|---|---|---|---|---|---|
| *Model* | *Treatment /Contrast* | *Year* | *Total* | *S. polaris* | *B. vivipara* | *Graminoids* |
| 1[a] | C | 2016 | 1.9 [1.24,2.57] | 1.33 [1.09,1.57] | 0.62 [0.38,0.86] | 0.46 [0.22,0.7] |
| 1[b] | I - C | 2016 | 0.04 [-0.23,0.32] | **0.27 [0.07,0.47]** | -0.09 [-0.29,0.1] | **-0.28 [-0.48,-0.09]** |
| | IW - C | 2016 | **-0.37 [-0.67,-0.06]** | **-0.5 [-0.71,-0.3]** | 0.15 [-0.05,0.36] | **-0.23 [-0.44,-0.03]** |
| | W - C | 2016 | -0.1 [-0.37,0.18] | **-0.54 [-0.74,-0.34]** | **0.23 [0.03,0.42]** | **0.26 [0.06,0.45]** |
| 2 [b] | I x W | 2016 | -0.31 [-0.8,0.17] | -0.23 [-0.57,0.1] | 0.02 [-0.32,0.35] | -0.21 [-0.54,0.13] |
| 1[a] | C | 2017 | 1.67 [1,2.34] | 0.87 [0.64,1.11] | 0.94 [0.7,1.18] | 0.13 [-0.1,0.37] |
| 1[b] | I - C | 2017 | -0.22 [-0.5,0.05] | 0.04 [-0.16,0.23] | **-0.24 [-0.43,-0.04]** | 0.01 [-0.19,0.2] |
| | IW - C | 2017 | 0.06 [-0.24,0.37] | -0.04 [-0.25,0.16] | 0.17 [-0.03,0.37] | 0.04 [-0.16,0.25] |
| | W - C | 2017 | -0.32 [-0.6,-0.05] | -0.42 [-0.62,-0.22] | -0.04 [-0.24,0.16] | 0.12 [-0.08,0.32] |
| 2 [b] | I x W | 2017 | **0.61 [0.13,1.1]** | **0.34 [0.01,0.68]** | **0.45 [0.11,0.78]** | -0.08 [-0.42,0.25] |

| | | | | | | |
|---|---|---|---|---|---|---|
| 1[a] | C | 2018 | 2.17 [1.5,2.84] | 1.44 [1.2,1.67] | 1.03 [0.79,1.26] | 0.44 [0.2,0.67] |
| 1[b] | I - C | 2018 | **-0.54 [-0.81,-0.27]** | **-0.42 [-0.62,-0.23]** | **-0.34 [-0.54,-0.15]** | -0.13 [-0.32,0.07] |
| | IW - C | 2018 | **-0.36 [-0.66,-0.06]** | **-0.42 [-0.62,-0.22]** | -0.1 [-0.3,0.1] | 0.01 [-0.2,0.21] |
| | W - C | 2018 | **-0.45 [-0.71,-0.2]** | **-0.66 [-0.85,-0.47]** | -0.16 [-0.35,0.02] | 0.19 [0,0.37] |
| 2[b] | I x W | 2018 | **0.63 [0.17,1.1]** | **0.67 [0.33,1]** | **0.41 [0.07,0.74]** | -0.06 [-0.39,0.28] |
| 1[a] | C | 2019 | 2.04 [1.38,2.71] | 1.09 [0.85,1.32] | 1.18 [0.94,1.41] | 0.4 [0.16,0.64] |
| 1[b] | I - C | 2019 | **-0.42 [-0.7,-0.15]** | -0.07 [-0.26,0.13] | **-0.36 [-0.55,-0.16]** | **-0.24 [-0.43,-0.04]** |
| | IW - C | 2019 | -0.23 [-0.53,0.08] | 0.03 [-0.17,0.24] | **-0.28 [-0.48,-0.07]** | -0.16 [-0.36,0.04] |
| | W - C | 2019 | **-0.33 [-0.6,-0.06]** | -0.35 [-0.54,-0.15] | **-0.34 [-0.53,-0.14]** | **0.2 [0,0.39]** |
| 2[b] | I x W | 2019 | **0.53 [0.04,1.01]** | **0.45 [0.11,0.78]** | **0.41 [0.08,0.75]** | -0.12 [-0.45,0.22] |
| 1[a] | C | 2020 | 2.69 [2.03,3.36] | 1.81 [1.57,2.04] | 1.01 [0.77,1.25] | 0.71 [0.48,0.95] |
| 1[b] | I - C | 2020 | 0.1 [-0.18,0.37] | **0.46 [0.26,0.65]** | **-0.4 [-0.6,-0.2]** | **-0.21 [-0.41,-0.01]** |
| | IW - C | 2020 | -0.21 [-0.51,0.1] | 0.1 [-0.1,0.31] | **-0.24 [-0.44,-0.03]** | **-0.28 [-0.49,-0.08]** |
| | W - C | 2020 | **-0.67 [-0.94,-0.4]** | **-1 [-1.19,-0.8]** | **-0.27 [-0.47,-0.07]** | **0.45 [0.25,0.65]** |
| 2[b] | I x W | 2020 | 0.37 [-0.12,0.85] | **0.64 [0.3,0.98]** | **0.43 [0.1,0.77]** | **-0.52 [-0.86,-0.19]** |
| 3[a] | C | 16-20 | 2.1 [1.43,2.76] | 1.31 [1.06,1.56] | 0.96 [0.71,1.2] | 0.43 [0.18,0.68] |
| 3[b] | I - C | 16-20 | **-0.23 [-0.46,0]** | 0.05 [-0.08,0.18] | **-0.3 [-0.43,-0.16]** | **-0.18 [-0.31,-0.05]** |
| | IW - C | 16-20 | -0.22 [-0.49,0.05] | **-0.17 [-0.31,-0.03]** | -0.06 [-0.2,0.08] | -0.13 [-0.27,0.02] |
| | W - C | 16-20 | **-0.35 [-0.58,-0.13]** | **-0.59 [-0.72,-0.46]** | -0.11 [-0.24,0.02] | 0.24 [0.11,0.38] |
| 4[b] | I x W | 16-20 | 0.36 [-0.05,0.77] | **0.37 [0.14,0.6]** | **0.35 [0.12,0.58]** | -0.19 [-0.42,0.04] |

| | | | | | | |
|---|---|---|---|---|---|---|
| 5 [a] | C | 16-18-19 | 2.09 [1.52,2.67] | 1.3 [1.11,1.49] | 0.96 [0.77,1.15] | 0.45 [0.26,0.64] |
| 5 [b] | I - C | 16-18-19 | **-0.44 [-0.71,-0.17]** | -0.11 [-0.25,0.04] | **-0.3 [-0.45,-0.16]** | **-0.26 [-0.4,-0.12]** |
| | IW - C | 16-18-19 | **-0.37 [-0.69,-0.05]** | **-0.31 [-0.46,-0.16]** | -0.09 [-0.24,0.06] | -0.14 [-0.3,0.01] |
| | W - C | 16-18-19 | **-0.32 [-0.59,-0.05]** | **-0.52 [-0.66,-0.38]** | -0.1 [-0.24,0.04] | 0.21 [0.07,0.35] |
| 6 [b] | I x W | 16-18-19 | 0.39 [-0.02,0.8] | **0.32 [0.07,0.56]** | **0.31 [0.06,0.56]** | -0.09 [-0.34,0.15] |
| 3 | Residuals | | 0.80 | 0.74 | - | - |
| | blocks/plots/sub-squares | | 0.11 | <0.01 | - | - |
| | blocks/plots | | 0.19 | 0.03 | - | - |
| | blocks | | 0.18 | 0.02 | - | - |
| | year | | 0.12 | 0.03 | - | - |
| | Observations | | 2880[c] | 8640[d] | - | - |

[a] Model prediction
[b] Model estimate
[c] 576 sub-squares (16 sub-squares replicated in 36 plots, plots are nested within 3 blocks), monitored over 5 years.
[d] Same as above but replicated for each three species groups.

Table S8. Temporal trends of Normalized Difference Vegetation Index (NDVI) metrics, relative abundance (Abund.) of vascular plants, specific leaf area (SLA) of *Salix polaris* and the community inflorescence count (Flower count). The fixed effects included treatment [as categories; *C* = control, *I* = icing, *W* = warming, *IW* = icing and warming combined] in interaction with year [as numeric] and the random intercept structure included plots nested within blocks (and sub-squares nested within plots for flower count). NDVI takes values from -1 to 1. max. = maximum; cum. = cumulative; Int. = intercept; std = standardized, sqrt = square-root. See Material and Methods for details.

| | *NDVI time of max. (day-of-year)* | *NDVI max.* | *NDVI cum. total* | *NDVI cum. start* | *NDVI cum. end* | *Abund. (log)* | *SLA (std and log)* | *Flower count (sqrt)* |
|---|---|---|---|---|---|---|---|---|
| Int. [*C*] | 198.82<br>(192.38 – 205.25) | 0.73<br>(0.70 – 0.76) | 0.7012<br>(0.6773 – 0.7251) | 0.71<br>(0.68 – 0.74) | 0.70<br>(0.68 – 0.72) | 3.22<br>(3.01 – 3.42) | 126.96<br>(23.16 – 230.76) | 1.51<br>(1.07 – 1.95) |
| [*I*] | 4.33<br>(-1.80 – 10.45) | 0.04<br>(0.01 – 0.07) | 0.0359<br>(0.0020 – 0.0697) | 0.02<br>(-0.02 – 0.06) | 0.05<br>(0.02 – 0.09) | 0.08<br>(-0.20 – 0.36) | -244.94<br>(-385.95 – -103.93) | -0.19<br>(-0.53 – 0.14) |
| [*IW*] | -1.21<br>(-7.33 – 4.92) | 0.05<br>(0.02 – 0.08) | 0.0598<br>(0.0259 – 0.0936) | 0.05<br>(0.01 – 0.09) | 0.06<br>(0.03 – 0.10) | 0.47<br>(0.20 – 0.74) | -58.87<br>(-201.22 – 83.47) | -0.23<br>(-0.60 – 0.14) |
| [*W*] | -2.96<br>(-9.09 – 3.16) | -0.00<br>(-0.04 – 0.03) | 0.0067<br>(-0.0272 – 0.0405) | 0.01<br>(-0.03 – 0.05) | 0.01<br>(-0.03 – 0.04) | 0.13<br>(-0.15 – 0.41) | -221.81<br>(-369.40 – -74.22) | -0.02<br>(-0.35 – 0.31) |
| year | -0.89<br>(-2.10 – 0.32) | 0.01<br>(0.01 – 0.01) | -0.0017<br>(-0.0070 – 0.0035) | -0.01<br>(-0.02 – -0.00) | 0.01<br>(0.00 – 0.01) | 0.05<br>(-0.00 – 0.10) | -0.06<br>(-0.11 – -0.01) | 0.20<br>(0.15 – 0.24) |
| [*I*] * year | 0.02<br>(-1.69 – 1.73) | **-0.01**<br>**(-0.02 – -0.00)** | -0.0084<br>(-0.0158 – -0.0010) | -0.01<br>(-0.02 – 0.00) | **-0.01**<br>**(-0.01 – -0.00)** | -0.02<br>(-0.10 – 0.06) | **0.12**<br>**(0.05 – 0.19)** | -0.01<br>(-0.07 – 0.06) |
| [*IW*] * year | 0.84<br>(-0.87 – 2.55) | **-0.01**<br>**(-0.02 – -0.01)** | -0.0128<br>(-0.0201 – -0.0054) | **-0.01**<br>**(-0.02 – -0.00)** | **-0.01**<br>**(-0.02 – -0.01)** | **-0.10**<br>**(-0.17 – -0.02)** | 0.03<br>(-0.04 – 0.10) | 0.00<br>(-0.06 – 0.07) |
| [*W*] * year | 0.11<br>(-1.60 – 1.82) | -0.00<br>(-0.01 – 0.01) | 0.0002<br>(-0.0071 – 0.0076) | 0.00<br>(-0.01 – 0.01) | -0.00<br>(-0.01 – 0.00) | -0.00<br>(-0.08 – 0.07) | **0.11**<br>**(0.04 – 0.18)** | **-0.11**<br>**(-0.18 – -0.05)** |
| Residuals | 29.5138 | 0.0003 | 0.0005 | 0.0010 | 0.0003 | 0.0317 | 0.8739 | 0.7887 |
| blocks/plots/sub-squares | | | | | | | | 0.0359 |
| blocks/plots | 0.1541 | 0.0008 | 0.0005 | 0.0004 | 0.0008 | 0.0396 | 0.1170 | 0.1199 |
| blocks | 17.3544 | 0.0001 | <0.0001 | <0.0001 | <0.0001 | 0.0003 | 0.0166 | 0.1045 |
| Observations | 144 | 144 | 144 | 144 | 144 | 144 | 3002 | 2880 |

Table S9. Summary statistic of the day-of-year a certain phenophase of *Salix polaris* is reached, illustrated in Figure S8. Estimated treatment effect sizes are reported in comparison to controls [C], on the log-scale with their associated 95% CIs. The response variable was the first day-of-year an inflorescence/leaf of a sub-square reached the phenophase. Two different generalized linear mixed-effect models were fitted to each phenophase: an additive model including 'treatment' [category] as a fixed effect and an interaction model between icing [*I*] and [*W*] treatments (binomial variables, *I* × *W*) as a fixed effect. The random intercept structure of the models were sub-squares [S] nested within plots [P], nested within blocks [B]. Phenophases correspond to Inflo vis. = inflorescence visible (female or male), Stigma vis = stigma visible, Stigma rec. = stigma receptive, Stigma sen. = stigma senesced, Seed disp. = seed dispersed, Anther vis. = anther visible, Pollen rel. = pollen released, Anther sen. = anther senesced, Unfurled = leaves unfurled, Expended = leaves fully expanded, Start sen. = leaves started senescing, Fully sen. = leaves fully senesced. σ2 = residuals. Bold numbers have their 95% CI excluding 0 (i.e., the C predicted mean).

| | **Inflo vis.** | **Stigma vis.** | **Stigma rec.** | **Stigma sen.** | **Seed disp.** | **Anther vis.** | **Pollen rel.** | **Anther sen.** | **Unfurled** | **Expended** | **Start sen.** | **Fully sen.** |
|---|---|---|---|---|---|---|---|---|---|---|---|---|
| Int. C | 5.06<br>(5.04 – 5.08) | 5.16<br>(5.13 – 5.18) | 5.19<br>(5.17 – 5.21) | 5.22<br>(5.21 – 5.24) | 5.39<br>(5.37 – 5.41) | 5.14<br>(5.13 – 5.16) | 5.17<br>(5.15 – 5.18) | 5.20<br>(5.18 – 5.22) | 5.07<br>(5.05 – 5.08) | 5.12<br>(5.11 – 5.14) | 5.30<br>(5.28 – 5.31) | 5.34<br>(5.32 – 5.35) |
| I | 0.01<br>(-0.02 – 0.04) | 0.01<br>(-0.02 – 0.05) | -0.01<br>(-0.04 – 0.03) | -0.00<br>(-0.03 – 0.02) | 0.02<br>(-0.01 – 0.05) | 0.02<br>(-0.01 – 0.04) | 0.01<br>(-0.01 – 0.04) | 0.00<br>(-0.02 – 0.02) | **0.03**<br>**(0.00 – 0.05)** | 0.01<br>(-0.01 – 0.03) | -0.00<br>(-0.02 – 0.02) | 0.00<br>(-0.01 – 0.02) |
| IW | 0.01<br>(-0.02 – 0.04) | -0.01<br>(-0.04 – 0.02) | **-0.03**<br>**(-0.05 – 0.00)** | -0.01<br>(-0.04 – 0.01) | **-0.05**<br>**(-0.08 – -0.03)** | **-0.03**<br>**(-0.05 – -0.00)** | **-0.03**<br>**(-0.05 – -0.01)** | -0.00<br>(-0.03 – 0.02) | **-0.02**<br>**(-0.04 – -0.00)** | **-0.03**<br>**(-0.04 – -0.01)** | 0.01<br>(-0.01 – 0.02) | -0.00<br>(-0.02 – 0.01) |
| W | 0.03<br>(-0.01 – 0.06) | 0.01<br>(-0.03 – 0.04) | 0.00<br>(-0.03 – 0.03) | -0.00<br>(-0.03 – 0.03) | -0.02<br>(-0.06 – 0.01) | -0.01<br>(-0.04 – 0.02) | -0.01<br>(-0.04 – 0.01) | -0.01<br>(-0.03 – 0.02) | -0.01<br>(-0.03 – 0.01) | -0.01<br>(-0.03 – 0.01) | 0.01<br>(-0.01 – 0.03) | 0.00<br>(-0.01 – 0.02) |
| I x W | -0.03<br>(-0.08 – 0.01) | -0.03<br>(-0.08 – 0.02) | -0.02<br>(-0.07 – 0.02) | -0.01<br>(-0.05 – 0.03) | **-0.05**<br>**(-0.09 – -0.01)** | **-0.03**<br>**(-0.07 – -0.00)** | -0.03<br>(-0.06 – 0.01) | 0.00<br>(-0.04 – 0.04) | **-0.04**<br>**(-0.07 – -0.01)** | **-0.02**<br>**(-0.05 – 0.00)** | -0.00<br>(-0.03 – 0.02) | -0.01<br>(-0.03 – 0.01) |
| $\sigma^2$ | 0.0063 | 0.0057 | 0.0056 | 0.0054 | 0.0046 | 0.0059 | 0.0057 | 0.0055 | 0.0063 | 0.0060 | 0.0050 | 0.0048 |
| B/P/S | 0.0000 | 0.0000 | 0.0000 | 0.0000 | <0.0001 | <0.0001 | <0.0001 | <0.0001 | <0.0001 | <0.0001 | <0.0001 | <0.0001 |
| B/P | 0.0006 | 0.0003 | 0.0003 | 0.0000 | <0.0001 | <0.0001 | <0.0001 | <0.0001 | 0.0001 | <0.0001 | <0.0001 | <0.0001 |
| B | 0.0000 | 0.0002 | 0.0001 | 0.0000 | 0.0001 | <0.0001 | <0.0001 | <0.0001 | <0.0001 | <0.0001 | 0.0001 | 0.0001 |
| Obs.[a] | 410 | 282 | 276 | 264 | 174 | 313 | 296 | 280 | 558 | 557 | 556 | 547 |

[a] Number of observations correspond to the number of sub-squares having at least one inflorescence/leaf in that particular phenophase. The total number of sub-squares is 576 (16 sub-squares in 36 plots).

Table S10. Summary statistic of the time spent in a certain phenophase (from the start of a phenophase to the next) in *Salix polaris*. Estimated treatment effect sizes are reported in comparison to controls [C], on the log-scale with their associated 95% CIs. For C, both the predicted means on the log-scale and rounded back-transformed values are reported. The generalized linear mixed-effect models had ‘treatment’ [category, *I* = icing, *IW* = icing x warming, *W* = warming] as a fixed effect and the random intercept structure included sub-squares [S] nested within plots [P], nested within blocks [B]. Inflo. Dev. = Inflorescence development from visible flowers (female or male) to stigmas/anthers visible; Stigma dev. = Stigma development from stigmas visible to stigmas receptive; Stigma rec. = from stigmas receptive to stigma senesced; Seed dev. = Seed development from stigma senesced to seed dispersal; Ather dev. = anthers development from anthers visible to pollen release; Pollen rel. = from pollen release to anthers senesced; Leaf dev. = leaves development from unfurling to being fully expanded; Expended = leaves fully expanded to starting to senesce; Leaf sen. = From leaves starting to senesce to fully senesced. σ2 = residuals. Bold numbers have their 95% CI excluding 0. The maximum number of sub-squares were 576 (16 sub-squares in 36 plots)

| | **Inflo. dev.** | **Stigma dev.** | **Stigma rec.** | **Seep dev.** | **Anther dev.** | **Pollen rel.** | **Leaf dev.** | **Expended** | **Leaf sen.** |
|---|---|---|---|---|---|---|---|---|---|
| Int. *C* | 2.29<br>(1.85 – 2.73)<br>~10 days | 0.93<br>(0.48 – 1.37)<br>~3 days | 1.43<br>(1.11 – 1.74)<br>~4 days | 3.38<br>(3.03 – 3.73)<br>~29 days | 0.75<br>(0.34 – 1.15)<br>~2 days | 1.68<br>(1.38 – 1.98)<br>~5 days | 2.15<br>(1.92 – 2.37)<br>~9 days | 3.46<br>(3.40 – 3.53)<br>~32 days | 1.88<br>(1.68 – 2.08)<br>~7 days |
| *I* | -0.33<br>(-1.01 – 0.36)<br>~7 days | **-0.74**<br>**(-1.44 – -0.03)**<br>**~1 day** | 0.03<br>(-0.42 – 0.48)<br>~4 days | 0.32<br>(-0.02 – 0.67)<br>~41 days | -0.03<br>(-0.58 – 0.52)<br>~2 days | **-0.48**<br>**(-0.90 – -0.07)**<br>**~3 days** | **-0.32**<br>**(-0.53 – -0.11)**<br>**~6 days** | -0.05<br>(-0.14 – 0.05)<br>~31 days | 0.10<br>(-0.18 – 0.39)<br>~7 days |
| *IW* | -0.44<br>(-1.06 – 0.19)<br>~6 days | -0.34<br>(-0.95 – 0.28)<br>~2 days | **0.41**<br>**(0.01 – 0.80)**<br>**~6 days** | **-0.35**<br>**(-0.64 – -0.05)**<br>**~21 days** | -0.05<br>(-0.61 – 0.51)<br>~2 days | **0.58**<br>**(0.17 – 1.00)**<br>**~10 days** | -0.05<br>(-0.25 – 0.16)<br>~8 days | **0.16**<br>**(0.07 – 0.25)**<br>**~38 days** | **-0.34**<br>**(-0.63 – -0.05)**<br>**~5 days** |
| *W* | **-0.85**<br>**(-1.53 – -0.17)**<br>**~4 days** | -0.01<br>(-0.69 – 0.66)<br>~3 days | 0.20<br>(-0.25 – 0.65)<br>~5 days | **-0.41**<br>**(-0.76 – -0.07)**<br>**~19 days** | -0.28<br>(-0.91 – 0.35)<br>~2 days | 0.17<br>(-0.27 – 0.62)<br>~6 days | -0.06<br>(-0.27 – 0.16)<br>~8 days | **0.10**<br>**(0.01 – 0.20)**<br>**~35 days** | -0.24<br>(-0.54 – 0.06)<br>~5 days |
| $\sigma^2$ | 1.27 | 2.27 | 1.01 | 0.35 | 1.78 | 0.39 | 0.21 | 0.03 | 0.56 |
| B/P/S | 1.13 | 1.85 | 0.82 | 0.32 | 1.37 | 0.23 | 0.09 | 0.00 | 0.40 |
| B/P | 0.27 | 0.14 | 0.05 | 0.03 | 0.12 | 0.13 | 0.03 | 0.01 | 0.06 |
| B | 0.00 | 0.00 | 0.01 | 0.06 | 0.00 | 0.00 | 0.02 | 0.00 | 0.00 |
| Obs. | 282 | 276 | 264 | 174 | 296 | 280 | 557 | 556 | 547 |

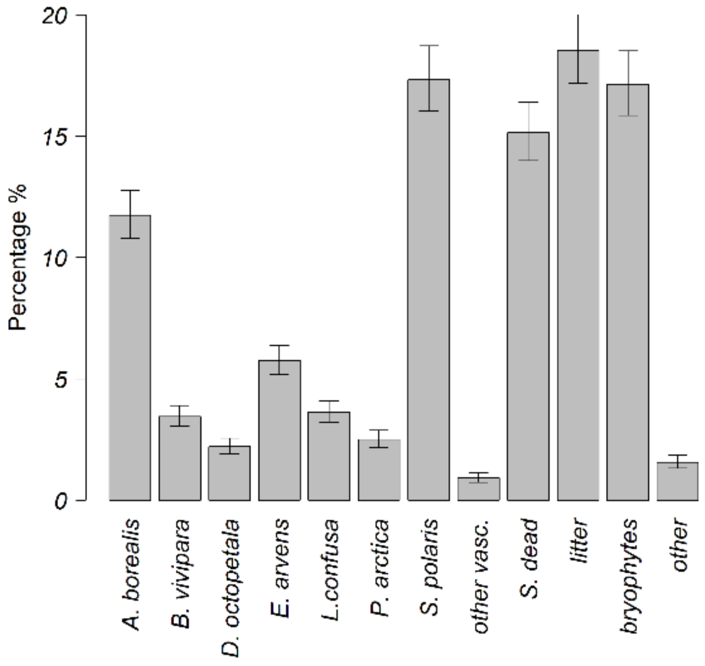


Figure S1. Mesic habitat species composition. Percentage of species or species groups recorded during point intercept monitoring (i.e., proportion of hits for a given species of the total hits). Estimates and their 95% confidence intervals were obtained from generalized linear mixed-effects model (Poisson distribution) with species/groups as fixed-effect and a random intercept structure including plot nested within block and year (2016-2019, 3 blocks ,12 plots, 1584 observations).

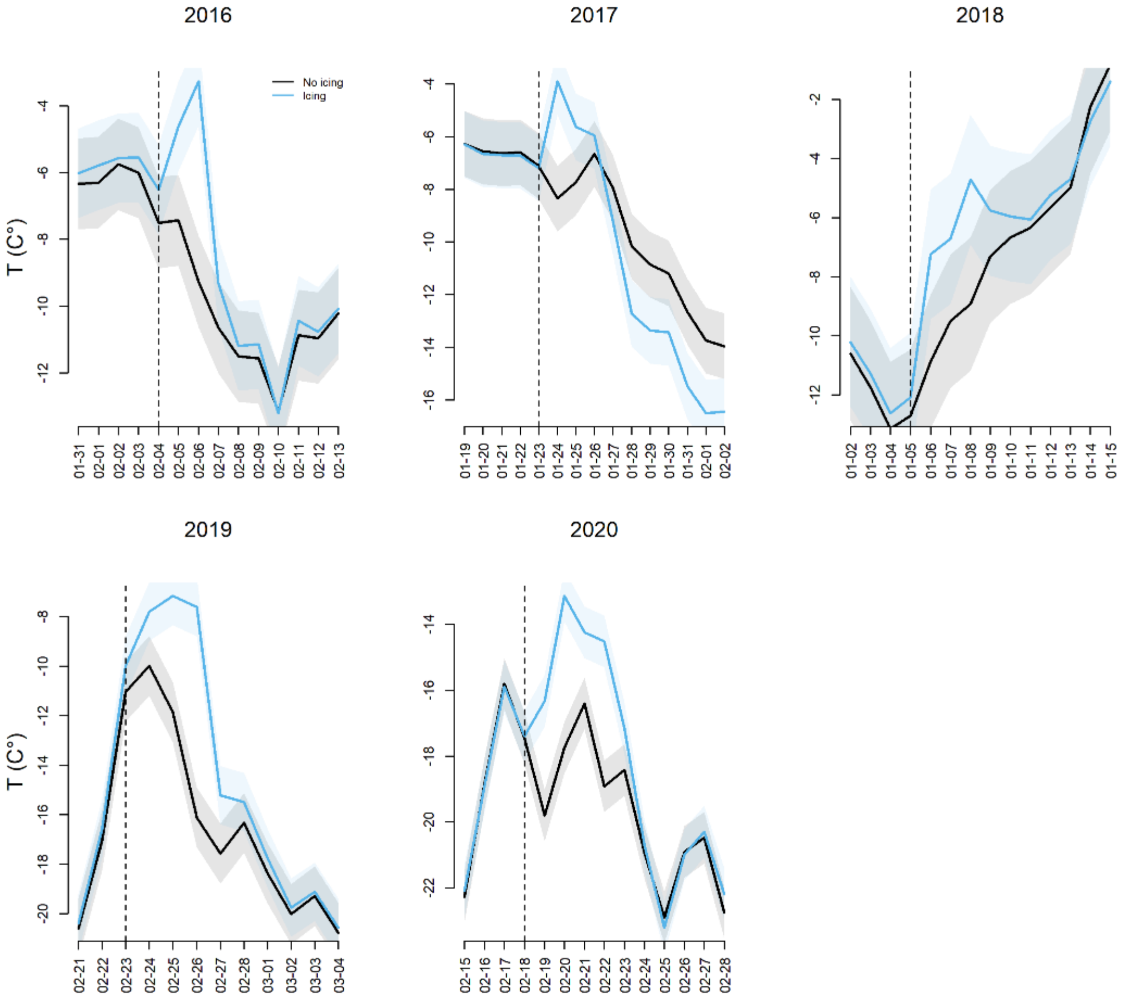


Figure S2. Daily estimated of sub-surface soil temperature (5 cm depth) in the period of the icing treatment simulation in January or February according to the year. Blue curves are the estimates of plots with simulated icing (i.e., icing and icing-warming), while the black curves did not receive any icing (i.e., control and warming). Vertical dashed line shows the day the icing treatment starts and lasts 2-3 days to obtain an ice layer of 13 cm thick. Linear mixed-effect models had treatment (i.e., icing yes/no) in interaction with day-of-year as fixed factors and the nested structure as random effect (i.e., plots within blocks). A separate model was fitted separately each year to reduce computation time. Shaded areas represent 95% confidence interval.

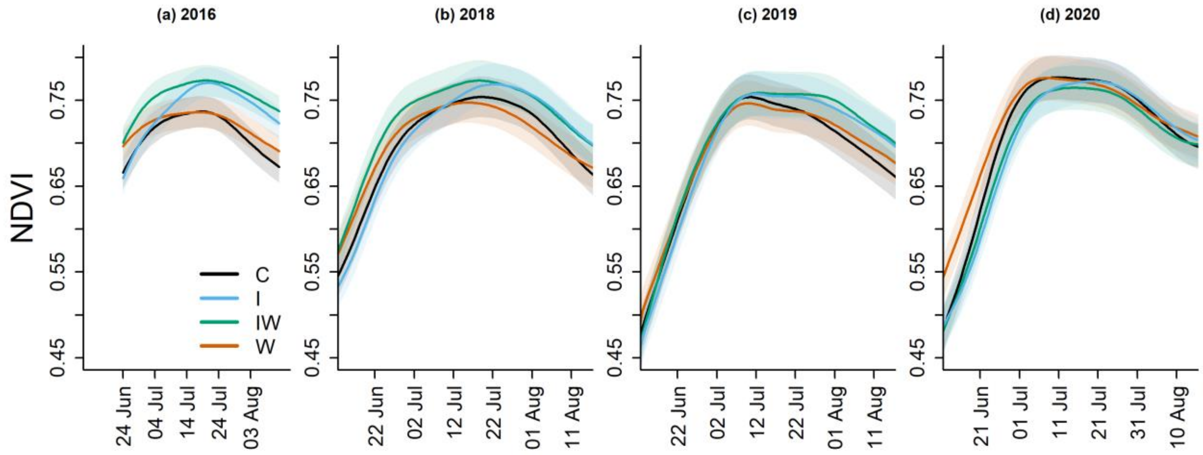


Figure S3. Estimated NDVI curves (with 95% CIs) across treatments and years. C = control, I = icing, IW = icing × warming, W = warming. The increased primary production of I and IW, in comparison to C, seemed to decrease over the years. However, this observation can be confounded with the natural fluctuations of ambient temperatures and timing of soil-thaw onset. For instance, the soil-thaw onset in spring 2020 occurred synchronously across blocks and plots (all control plots thawed over four days instead of e.g., over one month as in the year before, Table 1) and was followed by a heat wave in July. In that last year of the experiment, NDVI was very high across treatments, including controls, i.e., at the same level as in icing and icing × warming treatments in other years. This may reflect a contemporary 'threshold' in primary production of this community, beyond which it is possibly limited by nutrient availability.

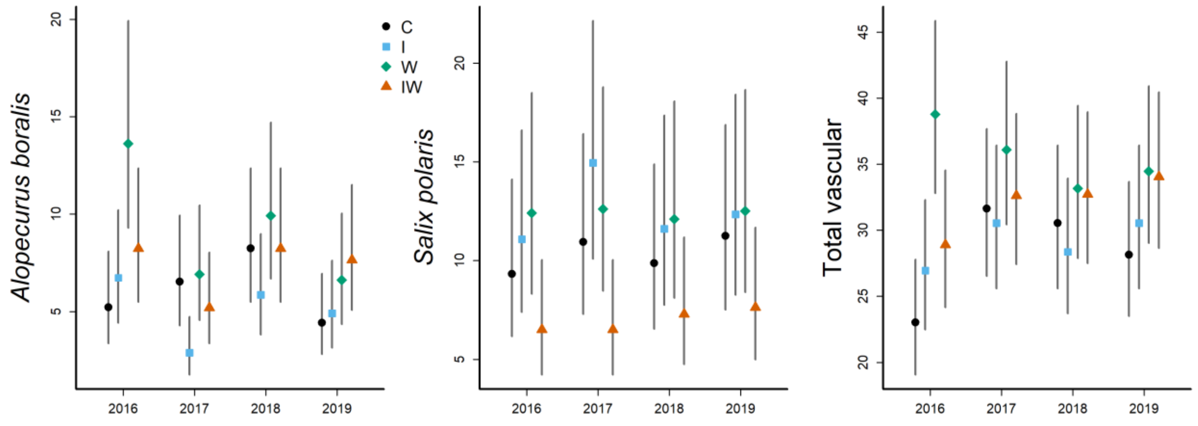


Figure S4. Annual relative abundance of two dominant vascular plants, (a) *Salix polaris* and (b) *Alopecurus borealis*, as well as (c) all vascular plants present (‘total vascular’). The y-axis represents the number of hits per plot (0.25 $m^2$). To estimate relative abundance, we used the point intercept methodology (Bråthen & Hagberg, 2004), with a 50 × 50 cm frame elevated above the canopy (~20 cm high) and with 25 evenly distributed points. The points were marked by crossings of double strings with the frame to give a 90º projection. At each point, a wooden pin of 3 mm diameter was lowered down onto the moss layer and all ‘hits’ of vascular plants were recorded. C = control, I = icing, IW = icing × warming, W = warming.

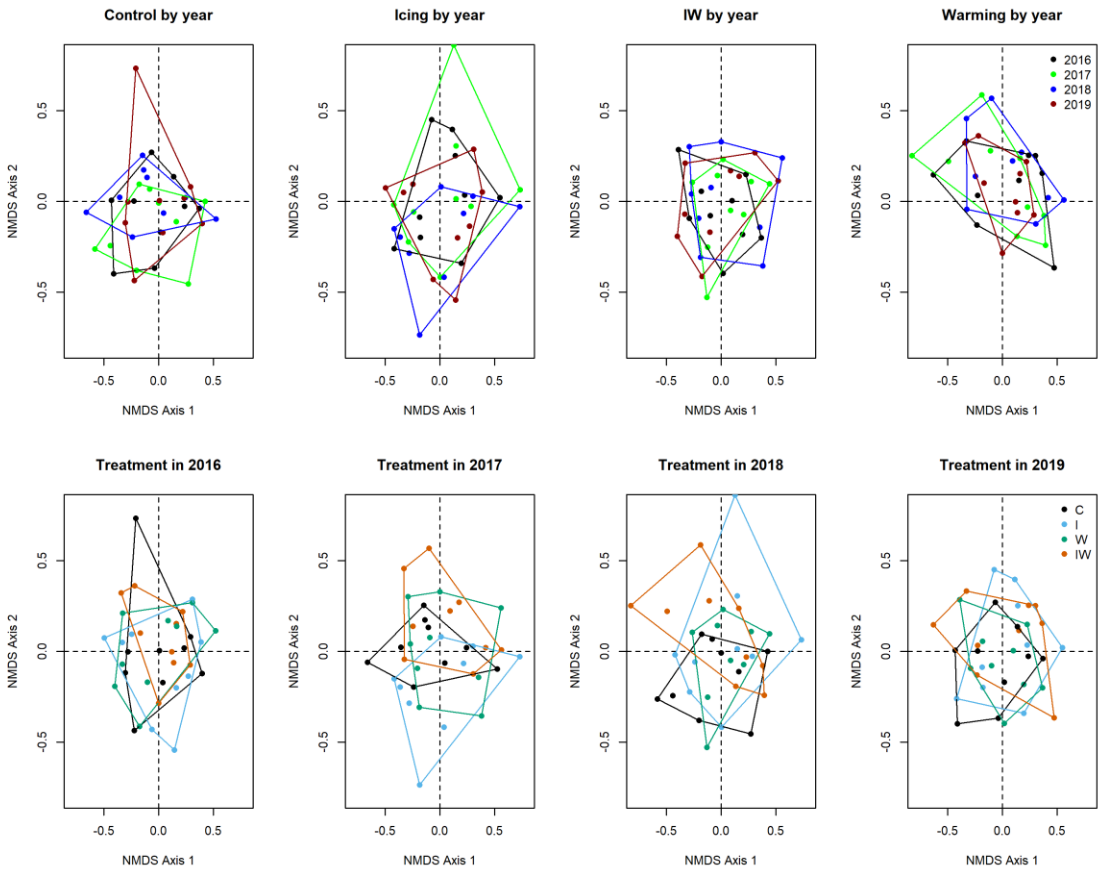


Figure S5. Non-metric multidimensional scaling (NMDS) ordination of the mesic community species composition. Points represent the plots' coordinates. The R-function 'metaMDS' in the vegan package was used. The large overlap of the ellipses demonstrates that the overall community compositions were similar across years and treatments. C = control, I = icing, IW = icing × warming, W = warming.

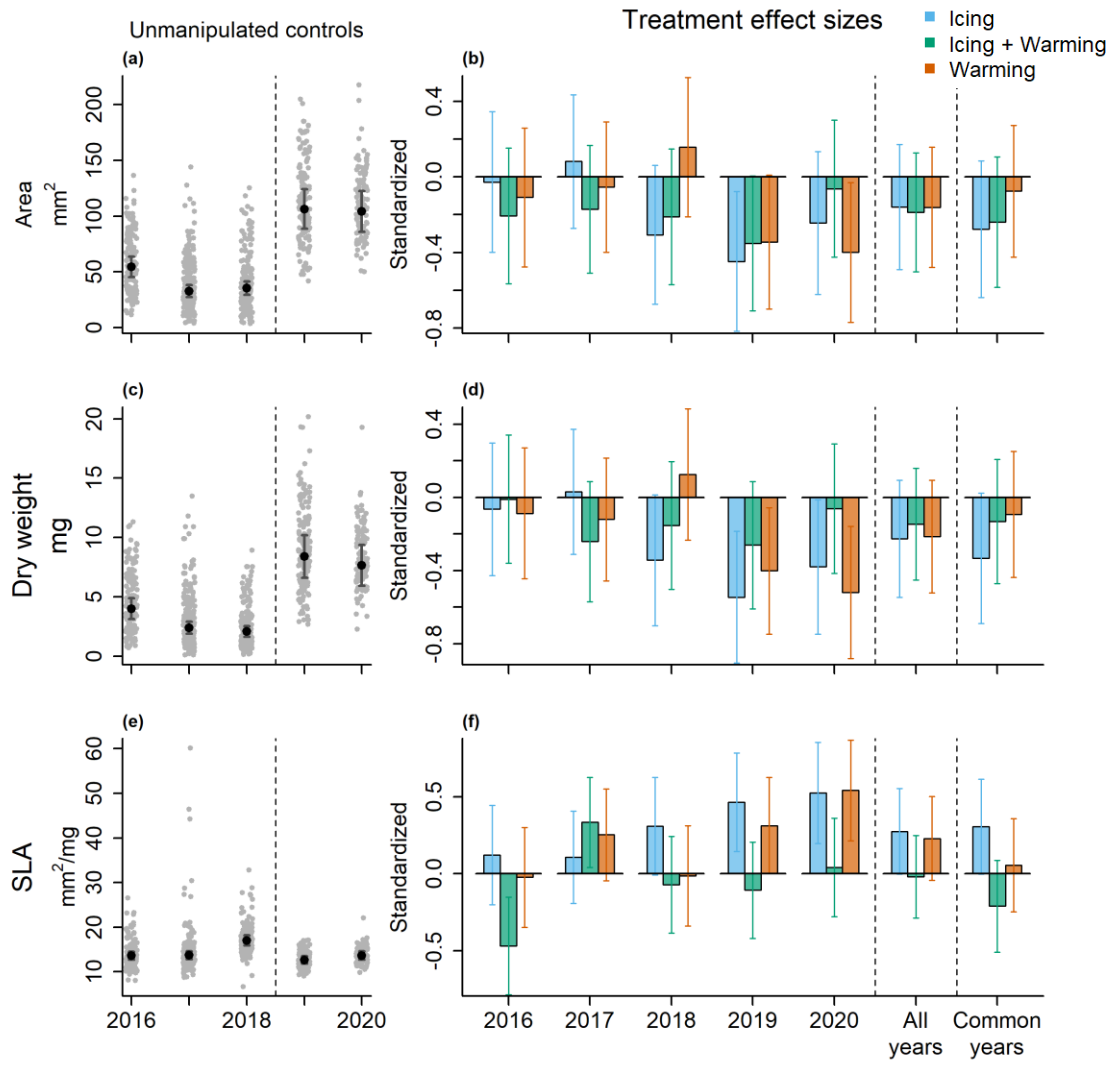


Figure S6. Effects of experimental treatments on leaf size traits of *Salis polaris* across years. The left panel (a, c, e) represents predicted means of control plots (black dots with their 95% confidence intervals) as well as the raw data (grey dots in the background), which were jittered for display purposes. The dashed vertical line mark when the sampling design shifted from random sampling of shoots (with all leaves nested within shoots measured), to sampling of the largest leaf per sub-plot. Therefore, the increase in leaf size and area observed in 2019 and 2022 is due to this change in sampling design. Model predictions and their CIs were back-transformed on the response scale prior presentation. On the right panel (b, d, f), we present the effect sizes and their 95% CIs for the effect of treatments displayed separately for each year and across years (year is used as a random-effect in the model, 'common years' included 2016, 2018 and 2019). The reference level of 0 corresponds to the predicted mean of controls (presented on the left panel). SLA = Specific leaf area, the ratio of leaf area to dry weight.

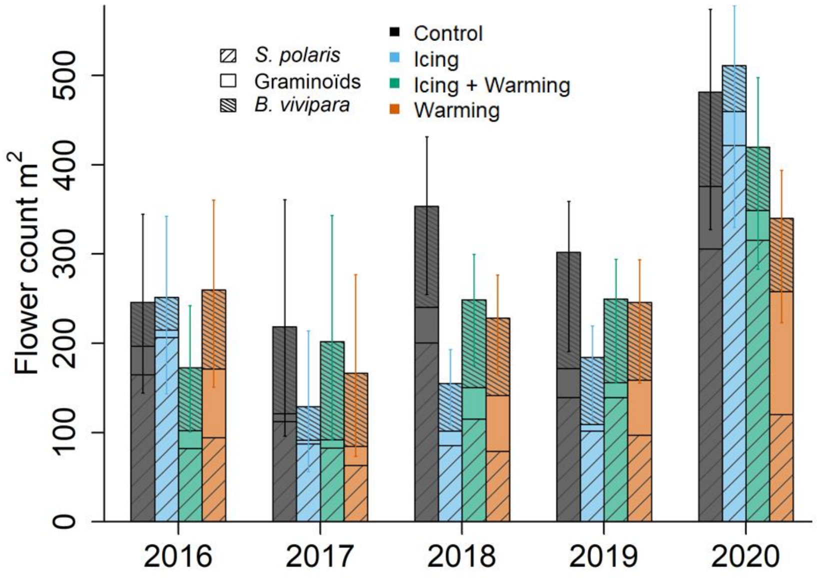


Figure S7 Flower/inflorescence counts per year, treatment and species/group. *S. polaris* and *B. vivipara* (combining reproductive shoots with inflorescence and/or bulbils) represent the species with the largest proportion of flowers in the community. The 'graminoids' group combines *A. borealis*, *L. confusa* and *P. arctica*. Predictions were obtained from generalized linear mixed-effects models, backtransformed from the square root-scale, with species × treatment × year included as fixed effect and a random intercept structure of plots nested within blocks. *C* = control, *I* = icing, *IW* = icing × warming, *W* = warming. The 95% CIs correspond to a model without the species distinction.

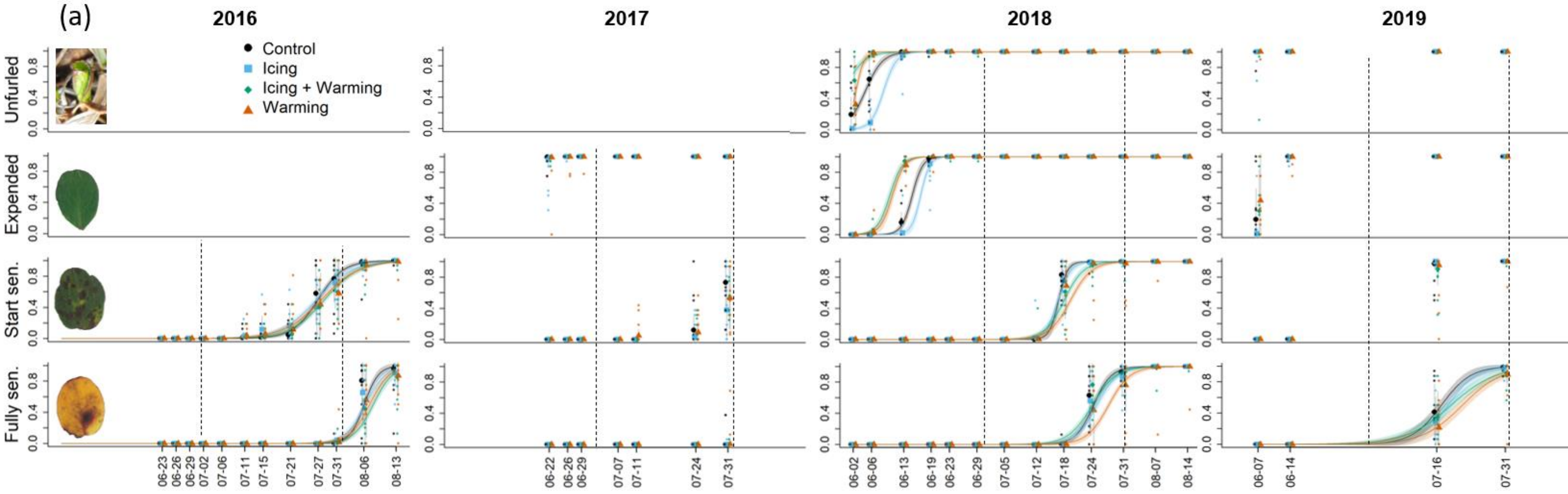

(a)
2016
2017
2018
2019
Control
Icing
Icing + Warming
Warming
Unfurled
Expended
Start sen.
Fully sen.
0.0
0.4
0.8
06-23
06-26
06-29
07-02
07-06
07-11
07-15
07-21
07-27
07-31
08-06
08-13
06-22
06-26
06-29
07-07
07-11
07-24
07-31
06-02
06-06
06-13
06-19
06-23
06-29
07-05
07-12
07-18
07-24
07-31
08-07
08-14
06-07
06-14
07-16
07-31

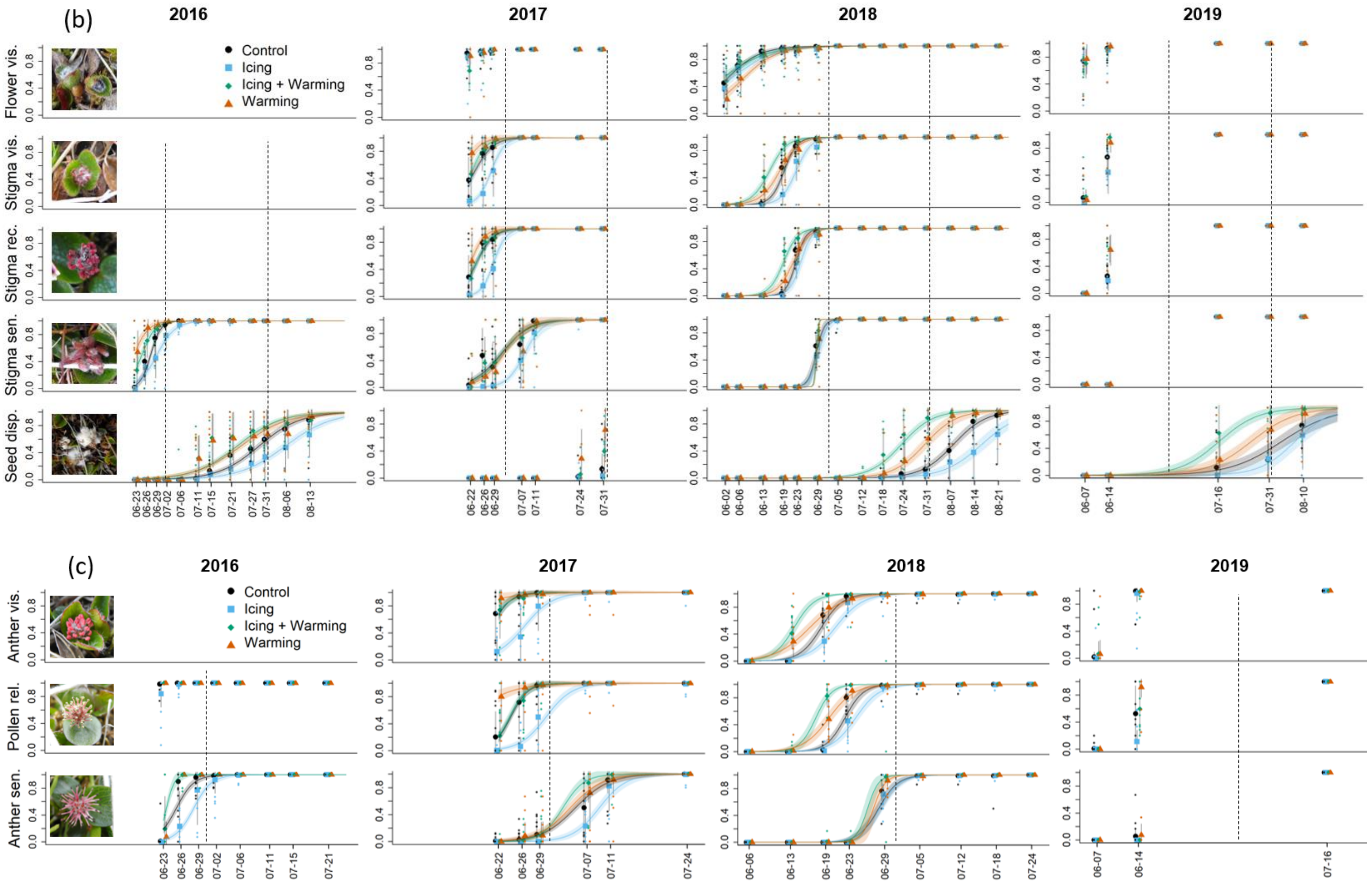

(b)
2016
2017
2018
2019
Control
Icing
Icing + Warming
Warming
Flower vis.
Stigma vis.
Stigma rec.
Stigma sen.
Seed disp.
(c)
Anther vis.
Pollen rel.
Anther sen.

Figure S8. Proportion of *Salix polaris* leaves (a), catkins (b) and male flowers (c) in a certain phenophase at each monitoring time-step from 2016 to 2019. The predicted mean proportion per treatment (main dots with associated 95% CI as grey bars) was estimated from a binomial distribution where the proportion of sub-squares in a certain phenophase within a plot was the response variable. The fixed effect was the interaction between 'treatment' and 'day-of-year' [as factor], fitted separately for each year. The replicated structure of the study design was accounted for by including 'plots' nested within 'blocks' as random effects. Similar models were fitted for continuous predictions across the field season (logistic curves), at the difference that the 'day-of-year' fixed effect was numeric (and not a factor). These curves were only calculated if the monitoring frequency was sufficient for reliable predictions. The data (i.e., proportion at the plot level) was represented by the small dots. The vertical dashed lines represent the month separation, on $1^{st}$ of July and $1^{st}$ of August. The phenophases were: (a) Unfurled = leaves unfurling, Expended = leaves fully expanded, Start sen. = leaves start senescing, Fully sen. = leaves fully senesced; (b) Flower vis. = flower visible (both for catkins and male), Stigma vis. = stigma visible, Stigma rec. = stigma receptive to pollen, Stigma sen. = stigma senescing, Seed disp. = seed dispersal; (c) Anther vis. = anther visible, Pollen rel. = pollen released, and Anther sen. = anther senesced.

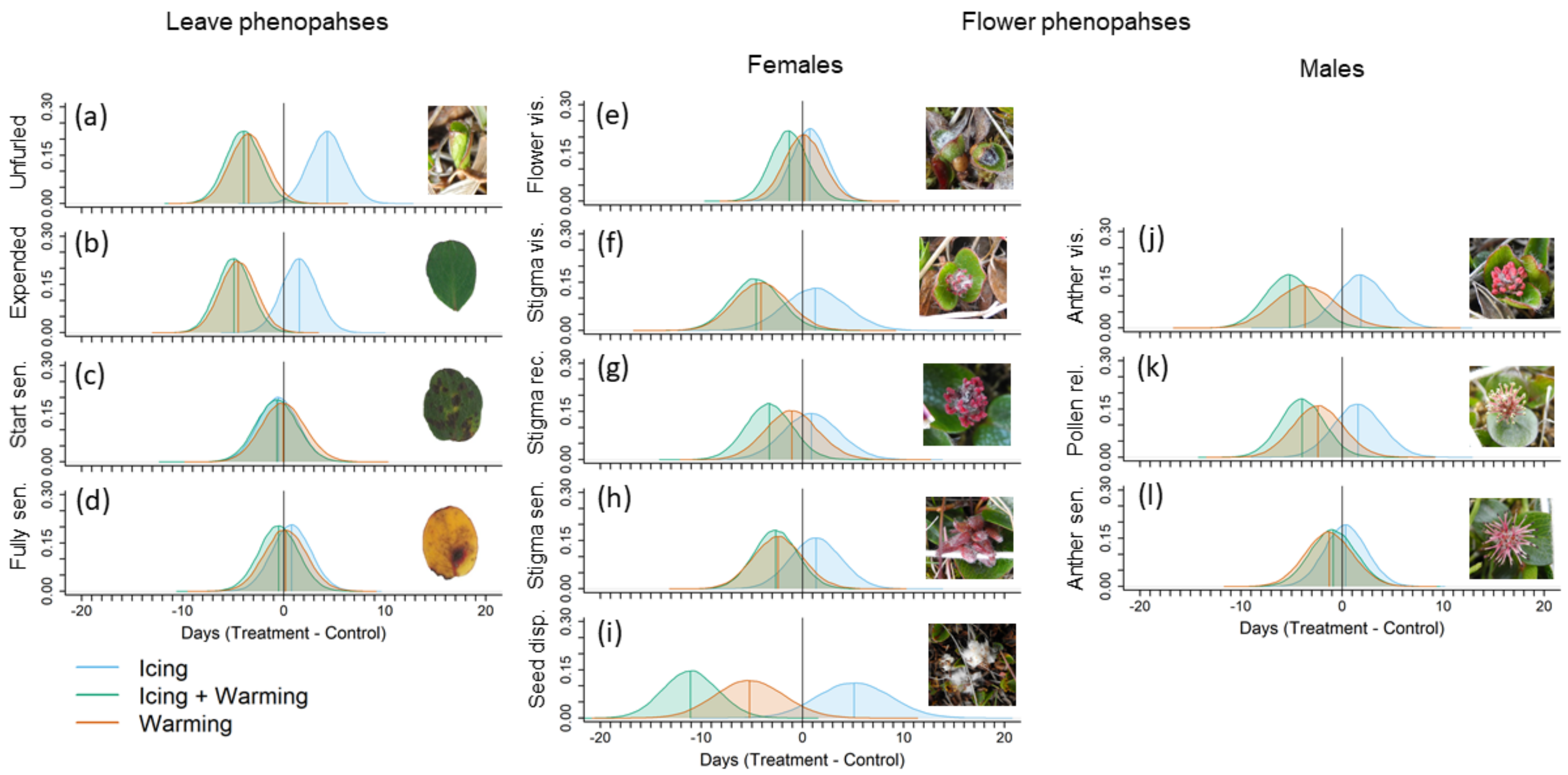


Figure S9. Differential distributions of the dates a certain phenophase of *Salix polaris* is reached (treatment – controls). Normal distributions were drawn from simulations with means and standard-errors obtained from generalized linear mixed-effect models following a Poisson distribution, accounting for the replication of sub-squares nested within plots, nested within blocks. The 0 vertical black line represents the intercept: the day-of-year (predicted mean) the control plots reached the phenophase being modelled. The coloured vertical lines represents the treatments' predicted means. Phenophases correspond to (a) leaves unfurled, (b) leaves fully expanded, (c) leaves started senescing, (d) leaves fully senesced, (e) flower visible (female or male), (f) stigma visible, (g) stigma receptive, (h) stigma senesced, (i) seed dispersed, (j) anther visible, (k) pollen released, and (l) anther senesced.
